# Negative design enables cell-free expression and folding of designed transmembrane β-Barrels

**DOI:** 10.1101/2025.09.16.676539

**Authors:** Giacomo Pedelli, Marvelous Chikerema, Andrei Sokolovskii, Tamas Lazar, Alexander V. Shkumatov, Alexander N. Volkov, Anastassia A. Vorobieva

**Affiliations:** Structural Biology Brussels, Vrije Universiteit Brussel, Brussels, Belgium; VUB-VIB Center for Structural Biology, Vlaams Instituut voor Biotechnologie (VIB), Brussels, Belgium; Jean Jeener NMR Centre, Vrije Universiteit Brussel (VUB), Pleinlaan 2, Brussels, Belgium

## Abstract

*De novo* design of membrane proteins (MPs) is a rapidly growing field with transformative potential for synthetic biology. Yet, progress has lagged behind that of soluble proteins, largely due to limited understanding of the fundamental principles governing MP folding, stability, and solubility—and their integration into computational models. Here, we use a cell-free expression system to bypass the cytotoxicity of failed or insoluble designs and investigate how sequence features influence the folding of synthetic transmembrane β-barrels (TMBs). We find that even small, idealized TMBs challenge classical protein design workflows: sequences optimized solely for thermodynamic stability misfold and aggregate, preventing membrane insertion. Instead of bulk hydrophobicity, aggregation and β-sheet propensity emerge as key determinants of membrane association. By designing better folding variants of a synthetic TMB, we demonstrate that suppressing aggregation-prone intermediates through local destabilization of β-strands (“negative design”) significantly improves folding efficiency. Strikingly, even substitutions typically considered highly destabilizing, such as prolines or polar threonines exposed to the bilayer core, can improve folding when strategically positioned, without significantly compromising thermodynamic stability. Based on these findings, we propose a framework for joint optimization of native stability and folding pathways for future MP and nanopore design.

**Significance Statement:** Membrane proteins are essential to biology and biotechnology, yet designing them from scratch remains challenging. Using synthetic transmembrane β-barrels and a cell-free expression system with lipid vesicles, we show that conventional design strategies focused solely on native state stabilization can lead to misfolding and aggregation. By incorporating “negative design” features—specific mutations that locally disrupt β-strand structure—we improve folding efficiency without compromising stability. Remarkably, a protein language model outperformed traditional energy-based methods in predicting these beneficial mutations. Our findings highlight the critical role of folding kinetics in membrane protein design and introduce new principles for engineering synthetic membrane proteins and β-barrel nanopores.

## Introduction

*De novo* protein design enables the generation of entirely new proteins, with no homology to natural sequences (1, 2). Over the past decade, the field has advanced from proof-of-concept demonstrations of custom protein folds and shapes (3, 4) to the generation of fully functional proteins for diverse applications (5–9). One approach to protein design — structure-based design — generates sequences based on 3D protein scaffolds that incorporate relevant structural or functional constraints (10). These methods optimize the amino acid sequence primarily for stability of the target native state, using physical energy optimization, statistical potential, or deep-learning models, while largely ignoring folding pathways and misfolded intermediates (11–13). Although this approach simplifies the complexity of protein folding, it has proven remarkably successful for designing idealized, water-soluble proteins, reinforcing the view that protein folding is, in general, governed by thermodynamics rather than kinetics (14).

This postulate has recently been challenged by efforts to design transmembrane β-barrels (TMBs), a class of β-sheet membrane proteins that span the lipid bilayer and form pore-like structures (15–18). Unlike soluble proteins, successful TMB design requires not only positive design (optimizing stabilizing interactions in the native state) but also negative design. In globular proteins, negative design typically prevents non-native states by adding repulsive contacts (19, 20). For TMBs, it instead involved introducing a small number of destabilizing features, such as disrupting the strict alternation of polar and hydrophobic residues along β-strands or incorporating disorder-promoting glycine or alanine residues. The precise mechanisms by which such subtle destabilization promotes successful folding remain unclear. Notably, sequences optimized solely through positive design consistently failed to express in *E. coli*, likely forming non-native cytotoxic species.

*In vitro* transcription-translation (IVTT) systems, such as PURExpress (21, 22), provide a powerful platform to examine the behavior of proteins that are toxic in cells (Figure 1A). When supplemented with lipid vesicles (Figure 1B), IVTT has been used to express and fold natural TMBs (23) and synthetic β-peptides (24). In this study, we use cell-free expression to dissect the role of negative design in TMB folding. We test synthetic sequences designed to adopt similar 8-stranded TMB structures but differ in biophysical properties, as well as variants carrying mutations that modulate stability and folding efficiency (Table S1). Our results suggest that negative design functions by suppressing premature folding or aggregation in aqueous solution. In contrast to conventional structure-based methods that have been broadly applied to the design of water-soluble proteins (10), we find that even small TMBs require a more nuanced framework that integrates both native stability and folding pathway constraints.

**Figure 1:**
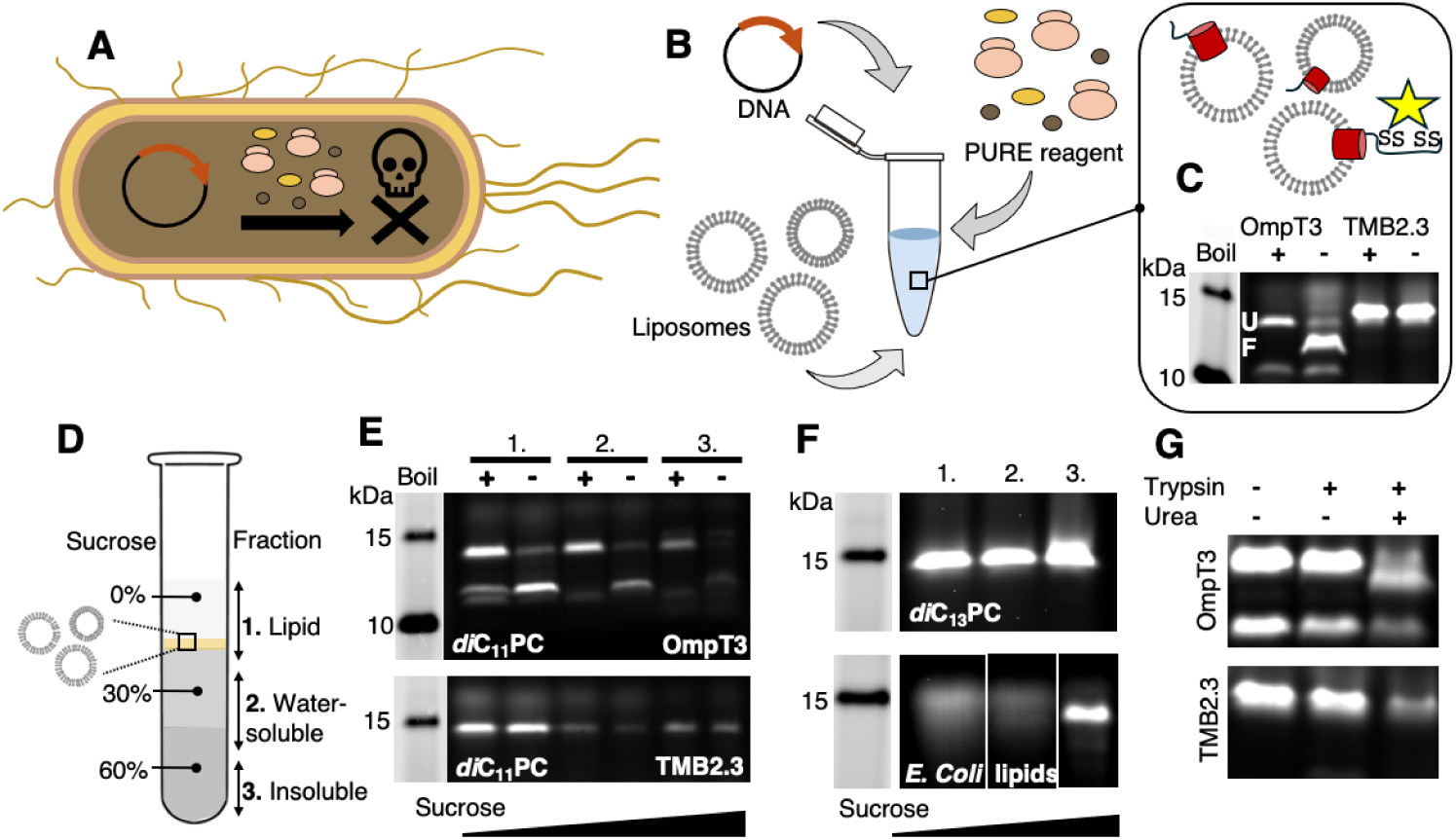
Cell-free protein expression of TMBs. (A) TMBs designed with the classic positive optimization paradigm do not express in E. coli. (B) Overview of cell-free expression. (C) Detection using a poly-cysteine reactive dye shows expression of OmpT3 (left, with characteristic heat-modifiable band) and TMB2.3 (right). (D) Overview of sucrose density gradient fractionation. (E) OmpT3 (top) and TMB2.3 (bottom) expressed with DUPC LUVs are enriched in the lipid-bound fraction. (F) Reduced interaction of TMB2.3 with lipid vesicles composed of longer aliphatic chain DMPC lipids (top) and E. coli polar lipids (bottom). (G) OmpT3 (top) and TMB2.3 (bottom) expressed with DUPC LUVs are resistant to protease digestion performed directly on the cell-free expression reaction.

## Results

### Expression and Membrane Insertion of TMBs in Cell-Free Systems

In biological membranes, TMBs require a dedicated BAM machinery to catalyze insertion and folding (25). By contrast, they can undergo spontaneous, thermodynamically driven folding into synthetic lipid vesicles. This unusual property, relative to most other membrane proteins, has enabled extensive thermodynamic studies of TMB folding from chaotropic denaturants into lipid bilayers (26–28). To validate folding during cell-free expression, we selected one native and one synthetic TMB based on their high folding efficiency (15). The first was the natural outer membrane protein OmpA, engineered with shortened extracellular loops to yield the more stable variant OmpT3. The second was the *de novo* designed 8-stranded TMB2.3, which has no detectable homology to natural TMBs and whose 3D structure has been validated by solution NMR. Because TMB folding efficiency depends on membrane properties such as hydrophobic thickness (26), we expressed both proteins in the presence of 1,2-diundecanoyl-sn-glycero-3-phosphocholine (DUPC, diC_11:0_PC) large unilamellar vesicles (LUVs; ∼ 100 nm diameter (Methods and Figure S1)). DUPC, a short-chain lipid, imposes a relatively low energy barrier to MP insertion (29). Both model TMBs were successfully expressed under these conditions, with a C-terminal poly-cysteine tag (CCPGCC) enabling in-gel detection using the fluorogenic biarsenical probe FLAsH-EDT_2_ (30) (Figure 1C). OmpT3 produced a characteristic heat-modifiable band on semi-native SDS-PAGE (like its parent OmpA), evidence of native folding for many TMBs (31), possibly resulting from interactions between the native state and SDS (32). By contrast, synthetic TMBs such as TMB2.3 typically do not exhibit heat-modifiable behavior (15), making the band-shift assay uninformative in this case (Figure 1C).

We further investigated the interaction of the TMBs with lipid membranes by separating insoluble (bottom fraction), water-soluble (middle fraction), and lipid-associated proteins (top fraction) using sucrose density gradient centrifugation (Figure 1D). As expected, most of OmpT3 was detected in the lipid fraction, where it retained its heat-modifiable behavior, indicating that it successfully integrates and folds into DUPC LUVs (Figure 1E, top). TMB2.3 was equally associated with the lipid fraction (Figure 1E, bottom). To distinguish integral membrane folding from mere surface adsorption, we tested TMB2.3 interaction with bilayers of varying thickness and opposing different kinetic barriers to folding (Figure 1F). Integral membrane folding would happen less efficiently in thicker bilayers (29), while adsorption to the membrane surface shouldn’t be affected by membrane thickness. Folding in the longer hydrocarbon chain lipid DMPC (1,2-dimyristoyl-sn-glycero-3-phosphocholine, diC_14:0_PC) resulted in reduced folding efficiency, as indicated by equal distribution of the protein across the lipid-associated and water-soluble fractions. When expressed in the presence of *E. coli* native lipids (where TMB folding requires the BAM machinery (25)), TMB2.3 aggregates instead of interacting with lipids, with a partition profile similar to that observed in the absence of lipid vesicles (Figure S2A). A similar effect of lipid composition was observed for OmpT3 (Figure S2B-D). Additional evidence for transmembrane folding in DUPC was provided by trypsin challenge, which demonstrated that OmpT3 and TMB2.3 were resistant to proteolysis (Figure 1G). Taken together, our results support integral membrane folding of both proteins.

We evaluated nanopore activity of IVTT-expressed TMBs using single-channel electrophysiology, where pore insertion into a lipid bilayer produces discrete current jumps that report on both pore number and structural state. To confirm that pores expressed in LUVs adopt the same conformation as those obtained by detergent refolding (a validated approach), we compared OmpT3 in both conditions. Refolded OmpT3 showed a symmetric base current of ∼±8 pA (Figure S3A). The same unitary conductance was observed for OmpT3 produced in LUVs (Figure S3B), although distributions were broader and multimodal, consistent with fusion of LUVs containing multiple pores (Figure S3A). These results indicate that both methods yield equivalent functional states. TMB2.3 expressed in LUVs also produced robust, voltage-dependent conductance steps (Figure S3C), confirming its folding into stable nanopores. Its higher conductance (∼±16 pA) relative to OmpT3 likely reflects the absence of side-chain gating seen in OmpA-derived pores (33, 34), as TMB2.3 forms a continuous transmembrane channel (Figure S4, channels detected with MOLEonline (35)). Together, these findings demonstrate that both natural and synthetic TMBs can fold into functional nanopores directly from cell-free expression, with conductance properties matching their expected architectures.

### Strong stabilization of the TMB folded state promotes aggregation

We next used the cell-free system to express TMB sequences designed with the classic native-state optimization approach: TMB0.1 and TMB1.1 (15). These designs were predicted by AlphaFold2 (36) to form stable β-barrel structures with high confidence (Figure S5). However, they had previously failed to express in *E. coli*. The first design tested, TMB0.1, was successfully expressed in the cell-free system, allowing us to investigate the origin of its problematic behavior. Density gradient ultracentrifugation revealed that the protein had a strong tendency to aggregate with all tested lipids (Figure 2A in DUPC, Figure S6 in *E. coli* lipids). TMB0.1 was designed with a highly charged lumen to promote pore hydration and was overall relatively hydrophilic (GRAVY hydrophobicity index: −0.541 (37)). To test whether its failure to interact with lipids was due to excessive hydrophilicity (by comparison, OmpT3 and TMB2.3 have GRAVY scores of −0.259 and −0.256, respectively), we expressed a second design, TMB1.1. This construct featured a polar but uncharged lumen, resulting in a more hydrophobic sequence (GRAVY index: 0.08). While TMB1.1 was also successfully expressed *in vitro*, it likewise failed to associate with membranes and instead aggregated (Figure 2B). Negative staining Electron Microscopy (nsEM) revealed that both designs formed dense aggregates co-localizing with clusters of ribosomes from the cell-free expression kit (Figure S7).

**Figure 2:**
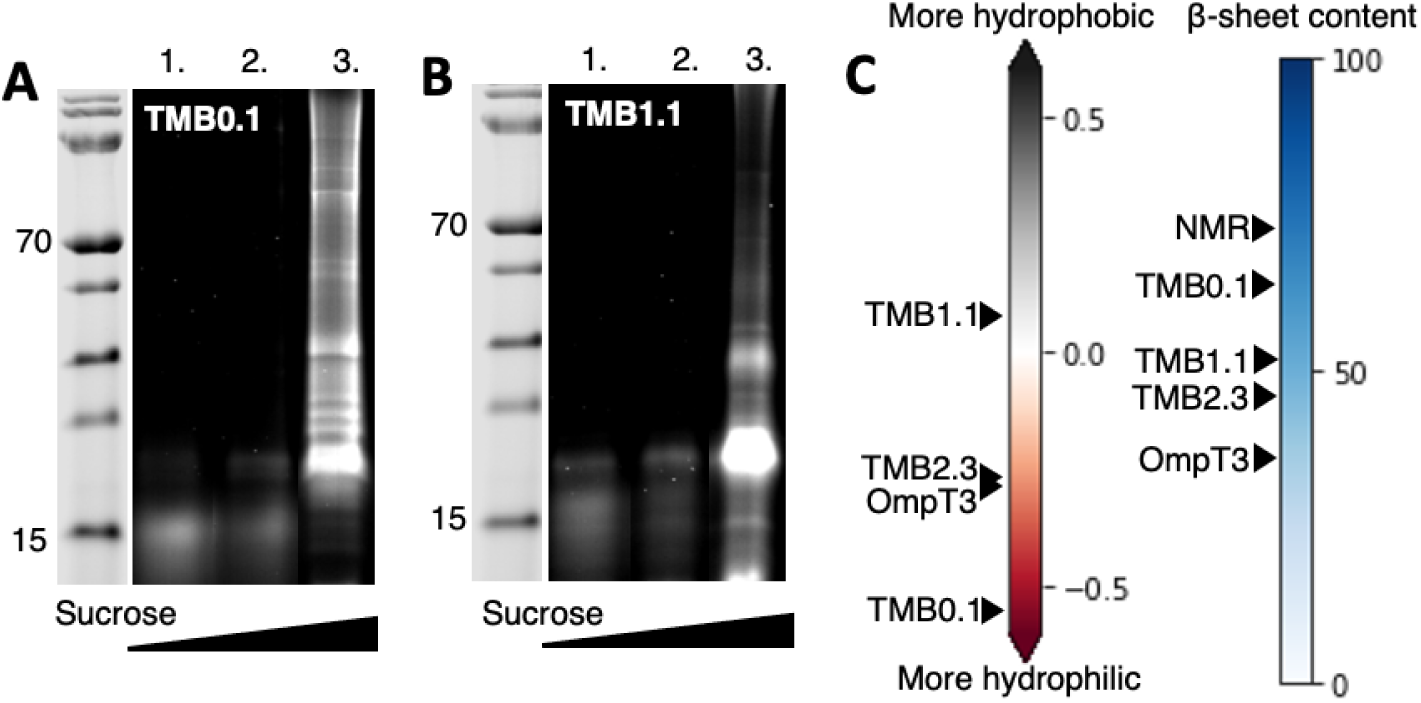
TMB0.1 and TMB1.1, designed with conventional optimization of native-state energy, aggregate due to high β-strand content. (A) TMB0.1 and (B) TMB1.1 aggregate after IVTT with DUPC LUVs, shown by enrichment in the bottom fraction (“3.”) after ultracentrifugation. (C) TMB0.1 and TMB1.1 hydropathy (GRAVY scale (37)) and β-strand propensity (percent of the total protein length, predicted by RaptorX (44)) compared to the synthetic TMB2.3 and the engineered OmpT3. NMR indicates the β-strand content of TMB2.3 determined from NMR chemical shifts (15). TMB designs with predicted β-strand content close to the experimentally determined content are more prone to aggregation.

These results indicate that strong optimization of the sequence for the native state can promote cross-β aggregation rather than assembly in the membrane. This likely explains the observed cytotoxicity of TMB0.1 and TMB1.1 (38, 39). While earlier studies linked efficient *in vitro* folding of native TMBs to high sequence hydrophilicity (40), our findings suggest that membrane association is not solely governed by bulk hydrophobicity. Despite having distinct hydrophobicities, both TMB0.1 and TMB1.1 aggregated instead of interacting with lipids (Figure 2C, Figure S6). We propose that high aggregation propensity stems from high β-strand content and structural order, which are more pronounced than in TMB2.3 and OmpT3 (Figure 2C). Notably, even natural TMBs vary in their tendency to aggregate into higher-order oligomers (28, 41, 42) or amyloid-like fibers (43), competing with proper membrane insertion and reducing folding efficiency over time (40). Our findings highlight this kinetic competition between aqueous aggregation and transmembrane folding as a key design constraint.

### Disorder-promoting mutations improve the folding efficiency of a synthetic TMB

Natural TMBs have evolved to achieve exceptional thermostability and folding efficiency (29, 45). By contrast, synthetic TMBs often display a wide range of stabilities and folding behaviors.

Understanding how these two properties are encoded within a single sequence is a critical step toward designing stable, functional MPs and nanopores that can be produced at high yield. We identified one 8-stranded design, TMB2.17 (15), which was highly stable but folded inefficiently, providing a model to probe the stability–folding tradeoff. Its β-barrel structure was previously validated by high-resolution X-ray crystallography, and it was the most stable design in the batch, with a folding free energy of 56 kJ mol⁻¹ in DUPC LUVs, compared to 38 kJ mol⁻¹ for TMB2.3 (15). Yet, TMB2.17 showed incomplete folding *in vitro*, partitioning between lipid-associated, water-soluble, and aggregated fractions after cell-free expression (Figure S8, Figure 3). On SDS-PAGE, it consistently produced two bands: one at the expected molecular weight and another faster-migrating band (Figure S9), likely reflecting partial folding or altered detergent binding, a phenomenon common to membrane proteins (46).

**Figure 3:**
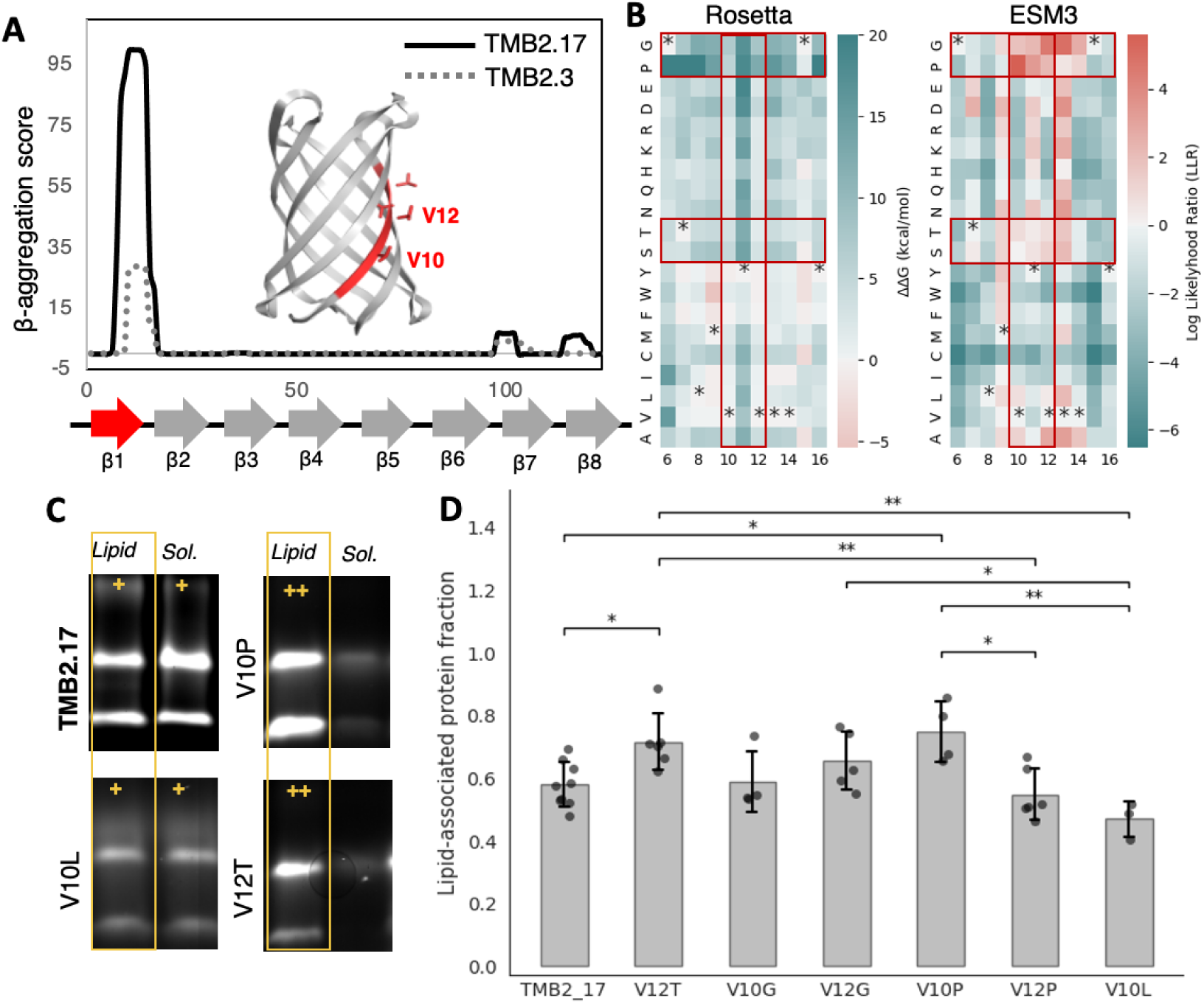
Mutations improving TMB2.17 folding. (A) Predicted β-aggregation hotspot in the first β-strand of TMB2.17. (B) Predicted effects of all possible single-point substitutions on β-strand1 on folding free energy (Rosetta, left) and evolutionary fitness (ESM3, right) (* indicates the WT per each position). (C) Representative SDS-PAGE gels showing partitioning of protein between the lipid-associated and water-soluble fractions after cell-free expression and density gradient separation. (D) Quantification of lipid association, calculated as Protein_lipid_ / (Protein_lipid_ + Protein_soluble_) fraction, increases from 58 % for wild-type TMB2.17 to 71-75 % for variants V12T and V10P. It remains at 55 % for V12P. (*p<0.05; **p<0.01).

We hypothesized that a local region in TMB2.17 retained a strong β-strand signature despite negative design, possibly interfering with proper folding. Aggregation propensity analysis with Tango (37) revealed a hotspot in the first β-strand, driven by clustered valines (V8, V10-12), with ∼3-fold higher aggregation score than the corresponding region in TMB2.3 (Figure 3A). *In silico* mutational scanning with Rosetta’s `cartesian_ddg` application (47) (Figure 3B) failed to identify mutations that would reduce the aggregation score without significantly destabilizing the native structure (Figure S10). We therefore turned to the ESM3 protein language model (48), which predicts evolutionary fitness rather than physical stability (Figure 3B). Surprisingly, several substitutions at V10 and V12 predicted as highly destabilizing by Rosetta were instead scored as beneficial by ESM3, including replacements with proline, glycine, or alanine—known to disrupt β-strands. Other predicted favorable mutations introduced polar residues (threonine or serine) into the otherwise hydrophobic barrel surface. These substitutions reduce aggregation propensity and increase predicted disorder of β-strand1 (IUPred3 (49), Table S1). Folding of TMBs *in vitro* likely involves concerted bilayer insertion and secondary structure formation (26, 27), with a lipid-embedded transition-state ensemble ordered on one side of the barrel and disordered on the other (28, 50). We reasoned that premature ordering of β-strand1—through secondary structure formation or aggregation—may interfere with assembly of this ensemble. To test this hypothesis, we constructed five variants (V12T, V10G, V12G, V10P, V12P) and expressed them *in vitro* with DUPC LUVs. Variants V10P and V12T showed significantly enhanced membrane association, whereas V10G, V12G, and V12P behaved similarly to wild-type TMB2.17 (Figure 3C-D). A control substitution, V10L, slightly reduced membrane association. Several β-strand-disrupting variants also decreased the intensity of the faster-migrating SDS-PAGE band, consistent with destabilization of a partially folded intermediate involving β-strand1 (Figure S11, Figure S12).

Together, these results show that residual β-aggregation hotspots can persist even in TMBs designed with negative design, hindering proper folding. Such hotspots were likely overlooked in earlier folding studies of TMB2.17 performed in 2 M urea, a condition that may destabilize the intermediate. Mutations that disrupt these regions improve folding efficiency. They were more accurately predicted with ESM3 (evolutionary fitness) than Rosetta (energy-based stability), underscoring the importance of kinetic factors in TMB folding that traditional stability models do not capture. Strikingly, the naturally evolved OmpT3 avoids such pitfalls, exhibiting lower aggregation and secondary structure propensities alongside higher predicted disorder than designed TMBs (Figure 2C, Table S1).

### TMB2.17 variants preserve native structure and high stability

Next, we evaluated the impact of folding-enhancing mutations on the structural integrity and stability of purified proteins refolded in detergent. The five TMB2.17 variants were expressed in *E. coli* as inclusion bodies and subsequently refolded in Fos-choline 12 (DPC) at twice the critical micelle concentration (CMC; see Methods). Size exclusion chromatography (SEC) revealed that all variants eluted as major peaks with retention volumes similar to the wild-type TMB2.17 (Figure S13). The most notable shift was observed for V12P, potentially indicating structural changes or altered detergent interactions. To assess secondary structure content, we performed far UV circular dichroism (CD) spectroscopy. All five variants had CD spectra characteristic of all-β proteins, nearly superimposable to that of wild-type TMB2.17. Deconvolution using ChiraKit (51) revealed a slight increase in stable β-sheet content in variants V12T (42 %) and V10P (41 %) compared to TMB2.17 (39 %) (Figure 4A, Figure S14A-B). In contrast, variants V10G and V12G showed a decrease in stable β-sheet content (33 and 34 %, respectively; Figure S14C-D), consistent with the introduction of disorder-promoting glycine. Variant V12P exhibited a small but distinct spectral shift at 230 nm (Figure S14E), suggesting a conformational change affecting the local environment of surface-exposed tryptophan residues. To examine tertiary structure, we used nuclear magnetic resonance (NMR) spectroscopy. ^1^H-^15^N HSQC spectra of the V12T variant displayed well-dispersed chemical shifts consistent with a folded β-barrel, resembling wild-type TMB2.17 (Figure 4C). TMBs exhibit distinctive ^1^H NMR signals between 8.5–10.5 ppm. One-dimensional ^1^H and ^1^H-^15^N HSQC NMR confirmed β-barrel-specific peaks in four variants—V10G, V12G, V12T, and V10P—supporting native-like folding (Figure S15A-D). In contrast, spectra of V12P suggested disrupted β-barrel formation (Figure S15A,E).

**Figure 4:**
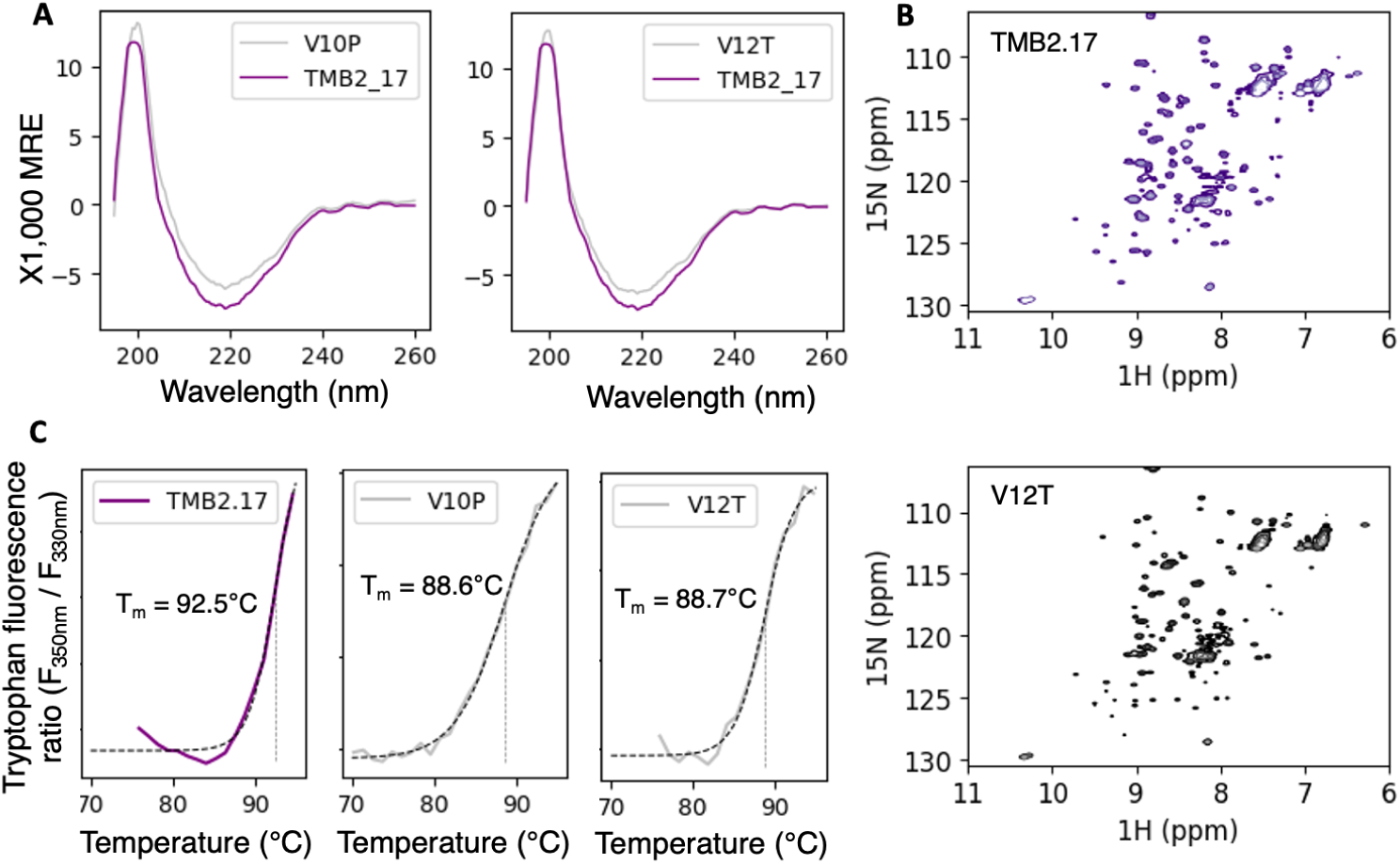
Biophysical characterization of TMB2.17 and its variants. (A) β-sheet characteristic circular dichroism spectra. Variants V12T and V10P have slightly higher β-sheet character (41 and 42 %, respectively) than TMB2.17 (39 %). (B) HSQC NMR spectra of TMB2.17 and V12T are characteristic of TMB proteins. (C) Thermal unfolding experiments in 3 M urea show cooperative two-state unfolding transitions. V12T and V10P mutations retain high thermostability.

Thermal stability in DPC micelles was evaluated by monitoring intrinsic tryptophan fluorescence during thermal denaturation. All proteins remained folded up to 95°C and required 3 M urea for detectable unfolding, following apparent two-state transitions (Figure 4B). Under these conditions, wild-type TMB2.17 had a melting temperature (Tm) of ∼92.5°C. The variants displayed a modest reduction in thermal stability, with ΔTm values ranging from 3.2 to 4.5°C. We further characterized thermostability across a range of urea concentrations (2.5-7 M) for TMB2.17, V12T, V12G, and V10P. All proteins unfolded between 6 and 7 M urea (Figure S16).

In summary, four ESM3-identified mutations predicted to be destabilizing based on traditional models preserved high thermodynamic stability and structural integrity of the TMB2.17 scaffold. Remarkably, V12T and V10P even enhanced folding efficiency in the cell-free system. The example of V12P highlights the difficulty of distinguishing between properly folded proteins and near-native, molten states based on standard biophysical methods alone.

## Conclusion

Our results highlight that designing transmembrane β-barrels (TMBs) requires more than maximizing thermodynamic stability. Positive design alone can drive misfolding and aggregation, whereas negative design delays premature folding in water and promotes correct membrane insertion (Figure 5). Yet even with negative design, residual β-aggregation hotspots can persist and hinder folding. In TMB2.17, such a hotspot in the first β-strand interfered with membrane integration; disrupting it with disorder-promoting mutations improved efficiency. This shows that folding bottlenecks can arise from local high-order “foldons,” and that selectively destabilizing them may be necessary to guide productive folding.

**Figure 5:**
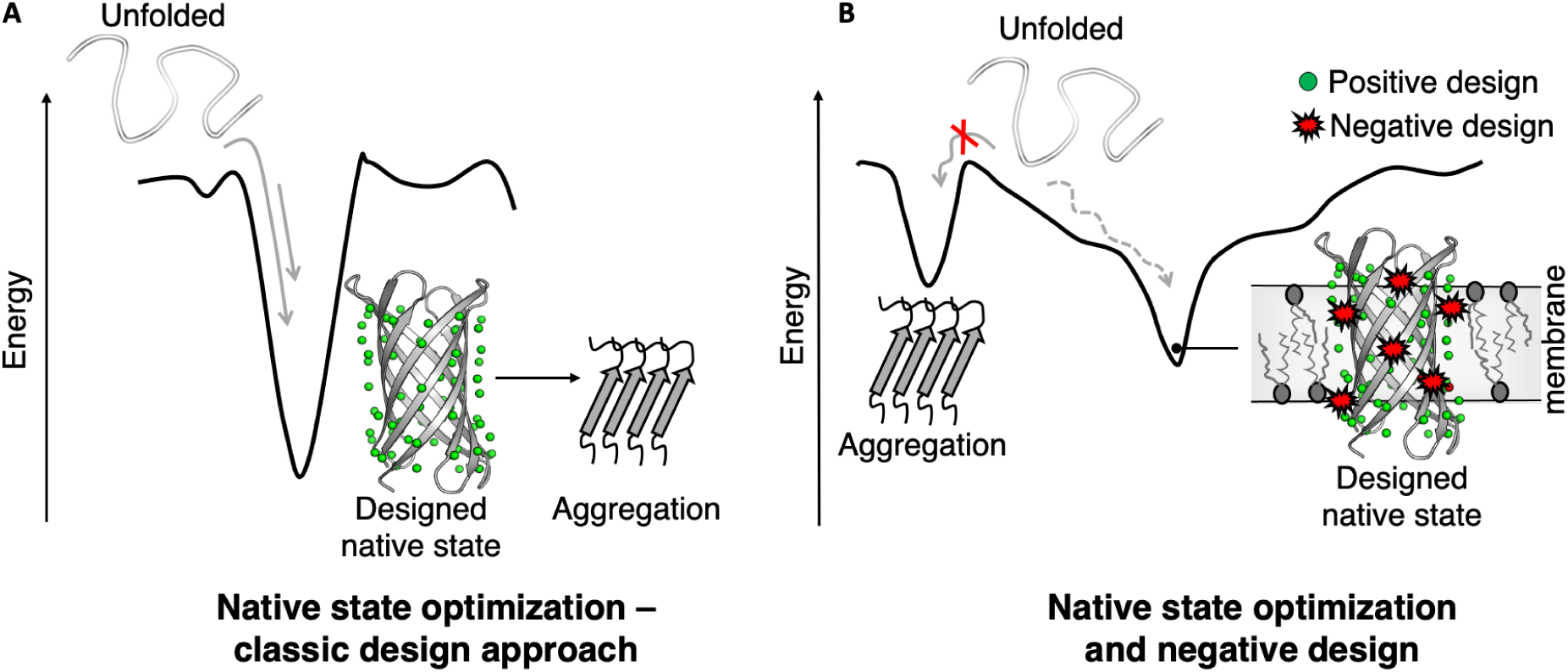
Proposed model of the interplay between native state stability and folding kinetics in designed TMBs. (A) Classic native state energy optimization leads to aggregation in water. (B) Negative design destabilizes local secondary structure elements, delaying early misfolding in water and promoting correct membrane insertion.

Notably, several folding-enhancing mutations (e.g., proline, glycine) had only modest effects on thermodynamic stability, suggesting that the membrane environment may buffer their destabilizing effects and pointing to gaps in our understanding of protein–lipid interactions. Although our experiments were conducted *in vitro*, folding kinetics are known to be critical for TMB folding *in vivo*, particularly in systems where unfolded precursors must traffic through cellular compartments before membrane insertion (e.g., TMBs crossing the periplasm in Gram-negative bacteria). Insights from synthetic, minimal TMBs may shed light on how natural TMBs achieve a balance between delayed folding during trafficking, overall high folding efficiency, and high stability after membrane insertion.

Together, these findings establish new design principles for synthetic β-barrels and underscore the need for computational methods that integrate both thermodynamic and kinetic considerations. Future strategies should combine physics-inspired models with machine learning approaches that model sequence fitness, such as ESM3. Such advances would be particularly important for the design of larger, more complex β-barrels, which have so far been challenging to design using traditional approaches due to computational limitations and the need to re-fit physics-based energy functions for these large proteins. Extending design capabilities to these larger nanopores could open the door to synthetic nanopores with pore sizes suitable for sequencing and molecular sensing applications. More broadly, these principles may also be relevant for soluble proteins, where folding kinetics and misfolded states are often overlooked in design, such as other membrane proteins and proteins whose folding depends on intermolecular interactions.

## Materials and Methods

### TMB expression in inclusion bodies and refolding in detergent

TMBs were produced in *E. coli* One Shot BL21 Star (DE3) cells, where, in the absence of a periplasmic-targeted signal peptide, they accumulate in inclusion bodies. Overnight cultures (25 µg/ml kanamycin, 0.5% glucose) were used to inoculate 50 ml of Studier autoinduction medium(52), which was incubated overnight at 37°C, 150 rpm shaking, to induce protein expression. The cells were harvested by centrifuging at 4000 rpm, 4°C, for 10 minutes on a TX-750 swinging bucket rotor (Thermo Fisher Scientific), resuspended in 30 ml of lysis buffer (50 mM Tris-HCl, pH 8, 40 mM EDTA, 150 mM NaCl), and disrupted by sonication. The lysate was mixed with 1% Brij-35 detergent and incubated for 30 minutes at 4°C, rotating at 20 rpm in an IKA loopster digital. The samples were then centrifuged for 50 minutes at 17,000 g at 4°C in a JA-14.50 rotor by Beckman Coulter Life Sciences. The pellet containing inclusion bodies was resuspending twice in wash buffer (10 mM Tris-HCl pH 8, 1 mM EDTA, 150 mM NaCl, 1% Brij-35), sonicated (Amplitude: 80% power; Pulse: 15 seconds ON, 15 seconds OFF; Time: 2 minutes sonication time), and centrifuged (17,000 g, 50 minutes, 4°C). The cycle was repeated twice without Brij-35 to remove residual detergent. The pellet was resuspended in 10 ml of denaturing buffer (6 M guanidine chloride, 25 mM Tris, pH 8) and centrifuged for 30 minutes at 10,000 rpm. The unfolded proteins were diluted to 80 μM in denaturing buffer and refolded by fast 20-fold dilution in 20 mM Tris-HCl, 150 mM NaCl, 0.1 % DPC as previously described (15).

### Liposomes preparation

25 mg of 1,2-diundecanoyl-sn-glycero-3-phosphocholine (DUPC), 1,2-dimyristoyl-sn-glycero-3-phosphocholine (DMPC), or *E. coli* Polar Lipid Extract were purchased from Avanti and dissolved in 1 ml of chloroform at room temperature (DUPC) or 37°C (DMPC and *E. coli* Polar Lipid Extract). 250 μl of the lipid solution were aliquoted into glass vials (6.25 mg/vial) and incubated in a vacuum overnight to allow chloroform evaporation. Lipid films were stored at −20°C. Liposomes were prepared for immediate usage by resuspension of the dry lipid films at 25 mg/ml in 10 mM Tris (pH 8.0), incubation on a rotator at room temperature for 30 minutes, and three freeze/thaw cycles in liquid N_2_. To obtain a monodispersed solution of large unilamellar vesicles (LUVs), the suspension was extruded with a minimum of 30 passages through 100 nm Track-Etch Membranes (Whatman^TM^) using a mini-extruder. To study size distributions of LUVs, extruded LUVs (25 mg/ml) were centrifuged to remove dust (10 minutes, 10,000 rpm) and analyzed using a DynaPro NanoStar DLS instrument from Wyatt Technology (10 acquisitions, acquisition time of 10 seconds, 25°C).

### Cell-free protein synthesis

For all cell-free protein synthesis experiments, the PurExpress^Ⓡ^ *in vitro* protein synthesis kit (New England Biolabs) was used according to the manufacturer’s instructions and for a total reaction volume of 30 µl. The reaction mix was supplemented with liposomes for a final concentration of 8 mg/ml of each tested lipid. The IVTT reaction was initiated by adding 200 ng of plasmid DNA (200 ng/µl), purified using phenol-chloroform extraction followed by ethanol precipitation to avoid residual nuclease contamination. Additionally, the reaction was supplemented with 20 units of RNAse inhibitor, Murine (NEB). *In vitro* protein expression was carried out for 3 hours at 37°C, and stopped by incubating the samples on ice for 10 minutes.

### Sucrose density gradient centrifugation

Sucrose solutions were prepared in 50 mM HEPES-KOH buffer at pH 7.6, 100 mM KCl, and 10 mM MgCl2 (assay buffer(22)). The gradient was prepared by depositing 50 µl of 60% sucrose in a 175 μl Ultra-Clear Tube (5 x 20 mm, Beckman Coulter), overlaid with 100 µl of 30% sucrose. Following IVTT, 30 µl of the PurExpress *in vitro* synthesis reaction were mixed with 20 µl of assay buffer and gently layered on top of the gradient. The tubes containing the gradients were centrifuged 3 hours at 250,000 g (45,000 rpm, Type 50.4 Ti Fixed-Angle Titanium Rotor from Beckman Coulter Life Sciences fitted with 5×20 mm adaptors (Beranek Laborgeräte)) to separate the lipid vesicles (floating at the interface between 30% sucrose and 50 mM HEPES-KOH buffer) and the unreacted components of the IVTT kit, pelletted to the bottom of the gradient. After centrifugation, 75 µl, 75 µl, and 50 µl were collected from the top to the bottom of the gradient to separate lipid-bound, water-soluble, and aggregated fractions. 12.5 µl of each fraction were analysed by SDS-PAGE (see below).

### SDS-PAGE and FLAsH-EDT_2_ labeling

Because high temperature treatment resulted in decreased dye efficiency, the samples were boiled (10 min, 95°C) in the Laemmli sample and cooled down to room temperature before labeling. The dye was dissolved in DMSO at 2mM stock concentration, and subsequently diluted in 10 mM Tris-HCl, pH 8. To label the proteins, they were mixed with Laemmli sample buffer, 25 mM TCEP, 0.1 µM FLAsH-EDT_2_ dye, and incubated for 30 min in the dark at room temperature. The samples were then analyzed by SDS-PAGE on 12% Tris-Tricine gels, running the experiment at 120 V for ∼90 min in the cold room to avoid overheating. The gels were visualized using a LI-COR Odyssey M machine (excitation: 685 nm; emission: 721-740 nm to visualize the ladder. Excitation: 488 nm; emission: 519-543 nm for the labeled proteins).

### Pulsed protease challenge

Trypsin digestion was performed directly on the cell-free reaction. 9 μl of samples were mixed with 0.1 mg/ml of Trypsin (lyophilized Recombinant Trypsin from Sigma-Aldrich resuspended in 1 mM HCl) and diluted up to 20 μL with H_2_O. For the control, the sample was prepared by adding 2 M of urea to destabilize the protein and allow trypsin digestion. The reactions were incubated at 37°C for 3 hours, and stopped by mixing the sample with Laemmli sample buffer and boiling for 10 minutes at 95°C.

### Electrophysiology

All-ion conductance measurements were performed using the Orbit mini instrument from Nanion Technologies on MECA 4 Recording Chips 150 μm. Lipid stock solutions of DPhPC (Di-Phytanoyl-Phosphatidyl-Choline) were freshly prepared before every recording experiment at a concentration of 5 mg/ml in dodecane. Lipid planar bilayers were painted on the four cavities to have a capacitance between 10-20 pF. Refolded proteins were diluted to a final concentration of ⁓0.25 mg/ml, and 0.2 μl of the proteins were added to the cis chamber of the chip containing 150 μl of KCl 1 M. 1 μl of cell-free expressed protein reaction was added to the cis chamber without further processing of the sample. Insertion pulses (100-200 mV) and 100 ms duration were used to help protein insertion. Spontaneous insertions of the protein were recorded at a temperature of 25°C. Signals were recorded using the Element Data Reader 4 software, setting a sampling rate of 10 kHz.

### Site-directed mutagenesis

The primers used for site-directed mutagenesis are listed in Table S1. All mutagenic PCRs were done using PrimeStar GxL polymerase according to the manufacturer’s instructions. Each reaction used 20 ng of template plasmid DNA. After amplification, the reaction mix was cleaned up (Wizard^Ⓡ^ SV Gel and PCR Clean-Up System by Promega) and incubated with DpnI restriction enzymes overnight at 37°C to digest unmutated, parental DNA. After DpnI inactivation (20 min, 80°C), the amplified plasmid was re-ligated using Gibson assembly (NEBuilder^Ⓡ^ HiFi DNA Assembly by New England Biolabs) in a total reaction volume of 10 µl. E. coli Top10 competent cells were transformed with 2 µl of the Gibson reaction mix.

### Negative stain electron microscopy

For negative stain data, 2 µl of the sample was applied on a glow discharged continuous carbon EM grid (FCF400-CU, EMS). After three short wash and blot steps with MilliQ water, the grids were immersed in 20 µl drops of 2% uranyl acetate for 10, 2, and 60 seconds with short blotting in between. Grids were visualized with a JEOL JEM-1400Plus electron microscope operating at 120 kV and equipped with a LaB6 cathode. Images were recorded on a CMOS TemCam-F416 camera (TVIPS, Germany) at a nominal magnification of 2,000 (pixel size 5.9 nm) and 60,000 (defocus of approximately 2.5 μm and a corresponding pixel size of 0.19 nm).

### Isotopic labeling for NMR experiments

A starter culture of *E. coli* One Shot BL21 Star (DE3) cells transformed with the expression plasmid was prepared in LB medium (with 0.5 % glucose and 25 µg/ml kanamycin) at an equal volume to the desired expression volume and grown overnight at 37°C, 200 rpm. Cells were harvested at 4,000 RPM, 4°C for 10 minutes and the pellet was gently resuspended in M9 minimal media (30mM Na2HPO4, 20mM KH2PO4, 10mM NaCl, 10mM NH4Cl, 0.2% (w/v) glucose, 1mM MgSO4, 0.1mM CaCl2, 0.01g/L biotin, 0.01g/L thiamin, 1x trace metals, 25 µg/ml kanamycin) with 1g/L ^15^N-NH4Cl. Cultures were grown at 37°C, 200 rpm, and protein expression was induced with 1mM IPTG at OD600 0.8-1.0. Cell cultures grown overnight at 22°C, 200 rpm, were harvested at 4,000 RPM, 4°C for 10 minutes. Supernatant was discarded, and the cell pellet was stored at −80°C or used immediately. Unfolded proteins were reconstituted in refolding buffer (see above), concentrated to ⁓200 μl using 15 ml 10K Amicon concentrators and dialyzed using Pur-A-Lyzer Maxi tubes (MWCO 6-8 kDa) with a ⁓100X NMR buffer (20 mM sodium phosphate buffer, 0.1% DPC, pH 6.8) excess, at 4°C for 3 hours. All samples were centrifuged for 2 minutes at 8000 rpm before the measurement. Unlabelled samples for ^1^H NMR were subject to additional purification step by SEC with a Superdex 200 Increase 10/300 GL by Cytivia and used at concentrations between 50-100 μM.

### NMR spectroscopy

Protein samples were prepared in 3mm NMR tubes and contained 20 mM sodium phosphate, 0.1 % DPC (pH 6.8) and 6 % D2O for the lock. All NMR spectra were acquired at 298 K on a Bruker Avance III HD 800 MHz spectrometer equipped with a TCI cryoprobe for enhanced sensitivity. 1D 1H spectra were recorded with 16 ppm spectral width, 4.7 ppm pivot, and 256 scans; water suppression was achieved by gradient-based excitation sculpting. 2D [1H, 15N} BEST-HSQC spectra were recorded with 800 and 256 points, 16 ppm and 35 ppm spectral widths, and 4.7 ppm and 117 ppm pivots in the direct and indirect dimensions, respectively, and 256 scans. All experiments were acquired and processed in Burker TopSpin 3.7.

### Thermostability assay

TMBs refolded in DPC micelles were purified by SEC (Superdex 200 3/10 Increase column, in refolding buffer: 25 mM Tris, 150 mM NaCl, 0.1 % DPC, pH 8.0). Samples were prepared at a concentration of 1 mg/ml in refolding buffer and supplemented with urea at concentrations ranging from 2.5 M to 7 M. 9 μl of sample were analyzed for tryptophan fluorescence using an Uncle instrument (Unchained Labs), ramping up the temperature 25°C to 95°C at a rate of 0.5°C/min (number of acquisitions: 5; acquisition time: 5 sec). A two-state sigmoidal transition was fitted to the recorded change of tryptophan fluorescence ratio (F_350nm_ / F_330nm_) to extract the T_m_.

### Circular dichroism

TMBs were refolded in DPC detergent micelles, purified through SEC, and analyzed at a concentration of 0.15 mg/ml. The CD spectrum was recorded on a MOS-500 instrument from BioLogic Science Instruments at a temperature of 25°C and with the following settings: wavelength range=190-260 nm; wavelength steps=0.5 nm; Acquisition period=0.5 sec; number of repeats=5. The spectra were analyzed with the ChiraKit tool (51).

### Structural modeling and bioinformatics

**AlphaFold2** (31) was used through the **ColabFold** implementation to assess TMB structure with the following settings: `recycles=3`; `single_sequence=True`; `template_PDB=False` and `AMBER_relaxation=True`. The **Rosetta** `cartesian_ddg` (43) protocol was used to evaluate the impact of point mutations on protein stability. The input structure (TMB2.17, PDB ID: 6X9Z) was prepared using Rosetta `FastRelax` in cartesian mode to ensure a physically realistic starting conformation. Three independent cartesian_ddg trajectories were calculated, averaged for each tested substitution and reported in a heat map. Aggregation-prone regions in protein sequences were identified using **TANGO** (37). Predictions were performed using the following conditions: pH 7.0, temperature 298 K, and ionic strength 0.02 M. A standalone version of **RaptorX-Property** was implemented for secondary structure prediction (40). All predictions were executed with no-profile mode, which does not rely on multiple sequence alignments. **ESM3** was used to predict mutational pseudo-likelihoods using the “esm3-sm-open-v1” model with openly available weights retrieved from HuggingFace (https://huggingface.co/). The transformer model was initialized in PyTorch using the ESM Python modules. The sequence was also tokenized using the “EsmSequenceTokenizer” module, then *in silico* site saturation mutagenesis was performed from the beginning to the end of the sequence with likelihoods converted to log-likelihood ratios (LLRs).

## Author contributions

A.A.V. designed research; G.P. and M.C. performed experiments. A.S. and T.L. ran simulations. A.V.S. performed nsEM. O.V. performed and analyzed NMR. G.P., M.C. and A.A.V. analyzed data. A.A.V. wrote the paper, with input from all authors.

## Acknowledgements

This work was supported by FWO (ERC runner-up grant FWOAL1092) and an ERC starting grant (PoreMaDNESS) to A.A.V. We thank the VIB Nanobody Core facility, Ema Romao in particular, for help with the Uncle instrument. We are also grateful to Prof. Gabe Rocklin and Prof. Peter Tompa for helpful feedback and suggestions.

## Data availability

The experimental and simulated datasets, the scripts and the Colab notebooks used for data analysis will be deposited on GitHub and archived in Zenodo at the time of the publication.

## Supplementary figures

**Figure S1.**
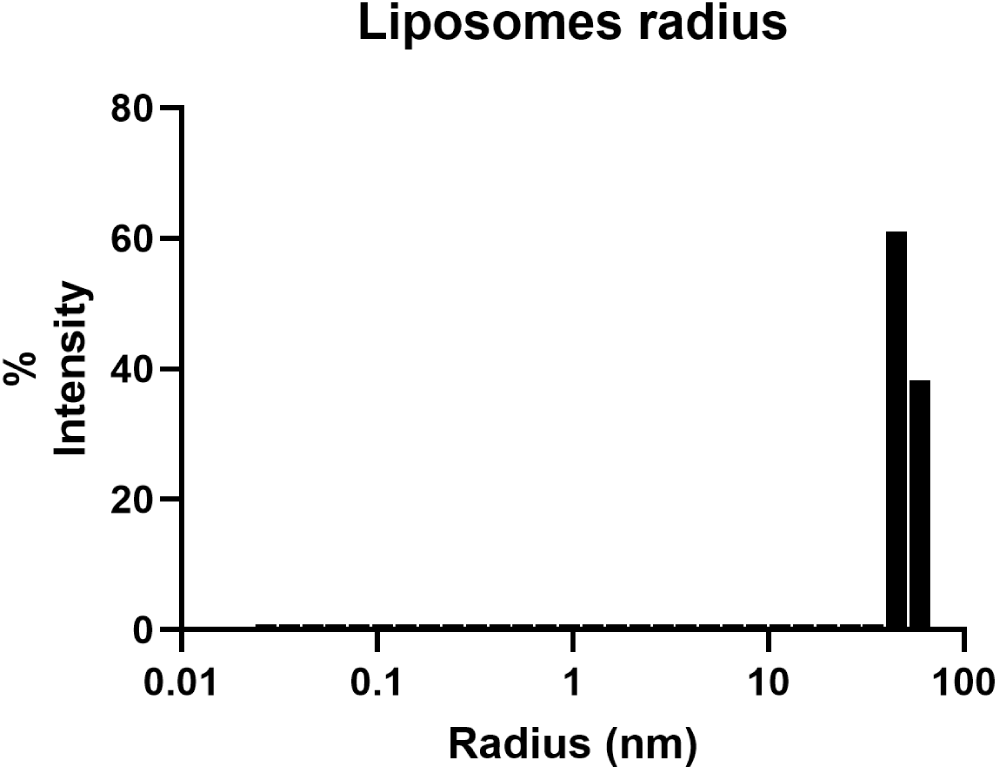
Representative Dynamic Light Scattering (DLS) analysis of extruded DUPC LUVs. The calculated diameter is 105.3 ± 2.5 nm (expected diameter ⁓100 nm) with a Polydispersity Index (PDI) of 0.19.

**Figure S2.**
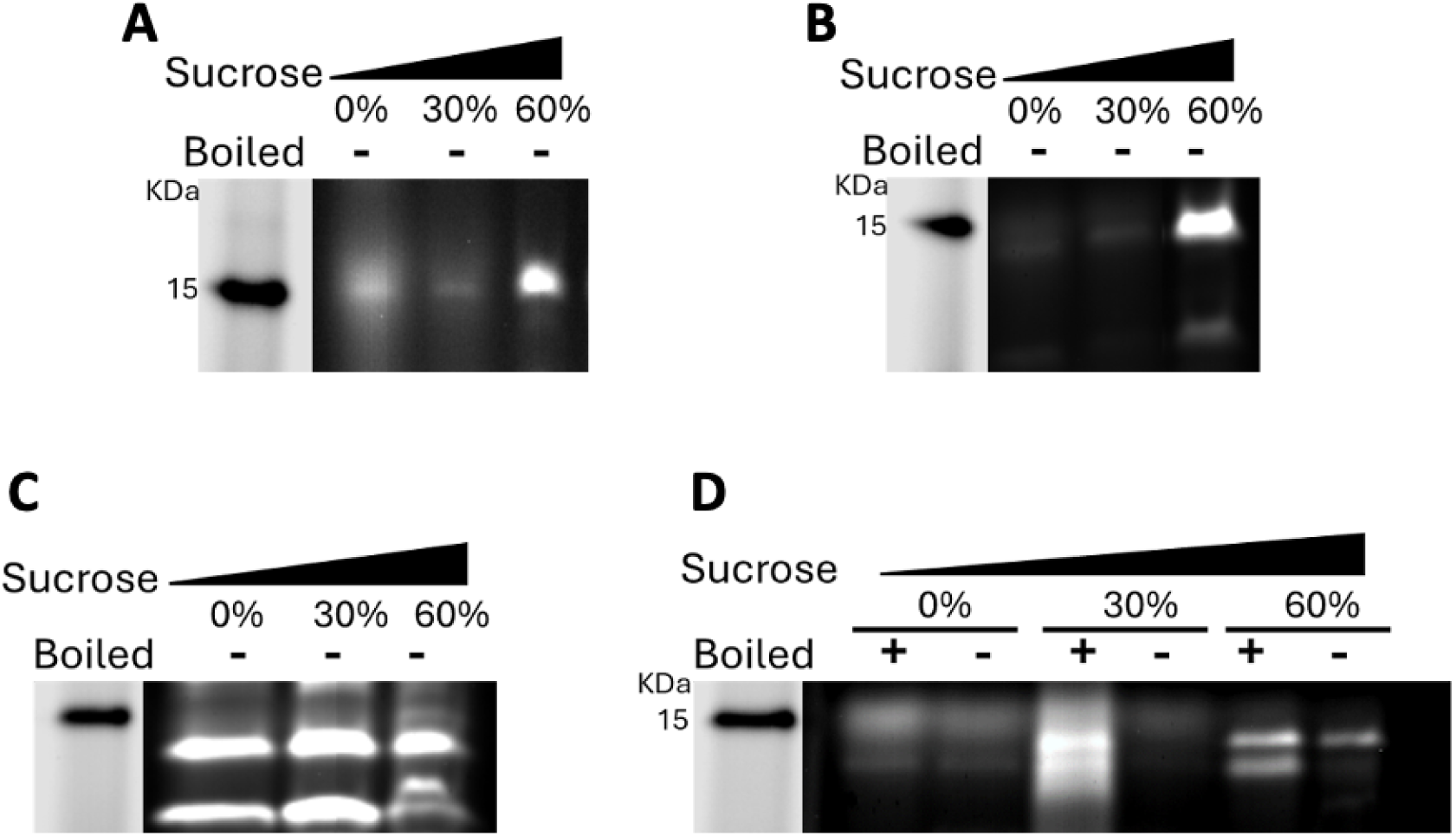
Partitioning profiles of TMB2.3 (A) and OmpT3 (B-D) after cell-free expression and sucrose density gradient ultracentrifugation in the absence of lipid (A-B); in the presence of DMPC LUVs (C); in the presence of E. coli Polar lipids LUVs (D).

**Figure S3:**
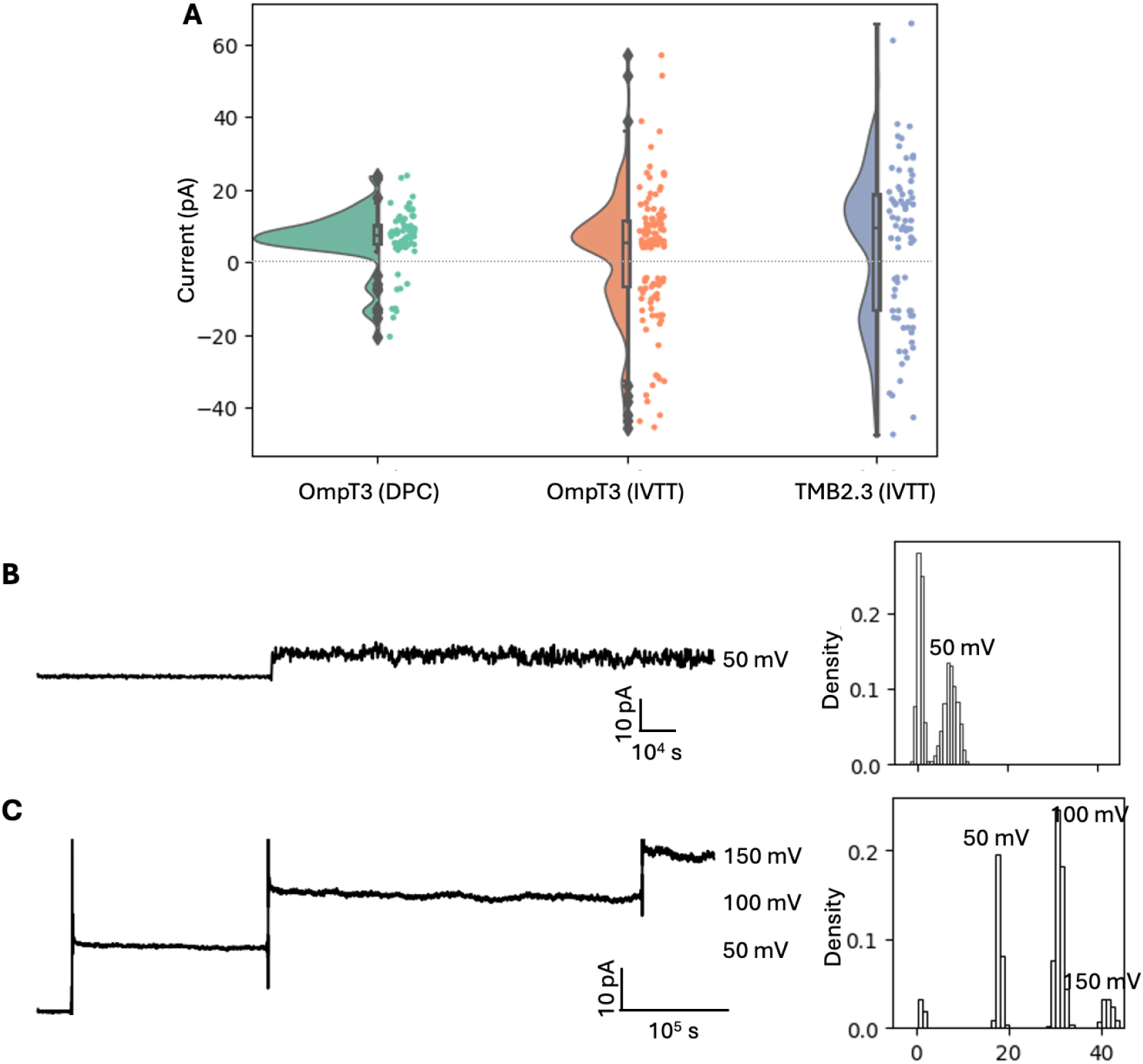
Single-channel electrophysiology. (A) Apparent base-pore current recorded for OmpT3 refolded in DPC micelles (N=71), OmpT3 IVTT-expressed in DMPC LUVs (N=122) and TMB2.3 IVTT-expressed in DMPC LUVs (N=77). Dots (right) show the conductances of independent recorded events; the densities of the dots are also shown (left). The box plots overlaid to density distributions show the median and interquartile range for each distribution. (B) Example single-channel insertion event of IVTT-expressed OmpT3 at 50 mV. (C) Example single-channel insertion event of IVTT-expressed TMB2.3 recorded at 50 mV, 100 mV and 150 mV.

**Figure S4:**
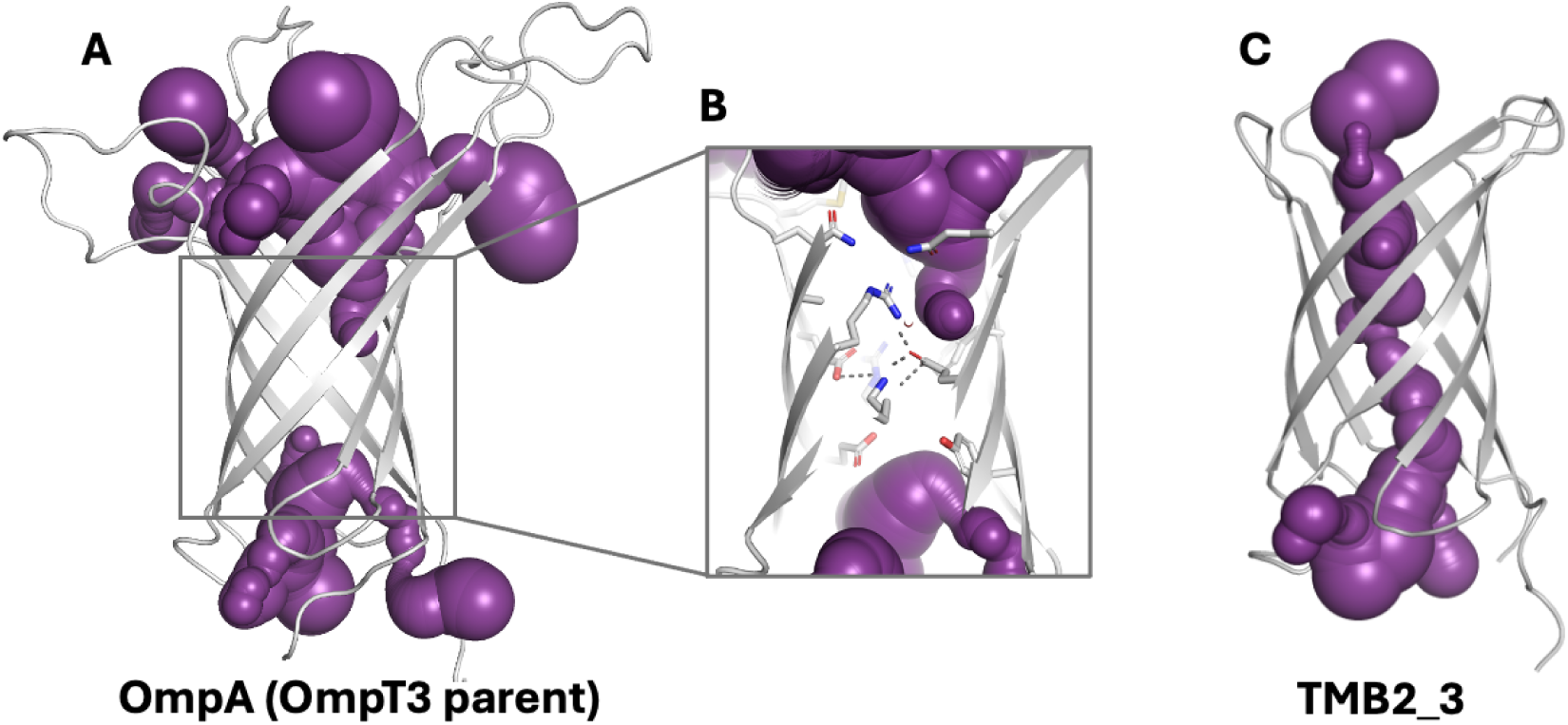
Channels detected using MOLEonline (purple) in (A) OmpA (parent of engineered OmpT3, PDB ID 2GE4); (B) OmpA with the side-chains forming the constriction that prevents a continuous transmembrane channel and (C) TMB2.3 (PDB ID 6X1K).

**Figure S5.**
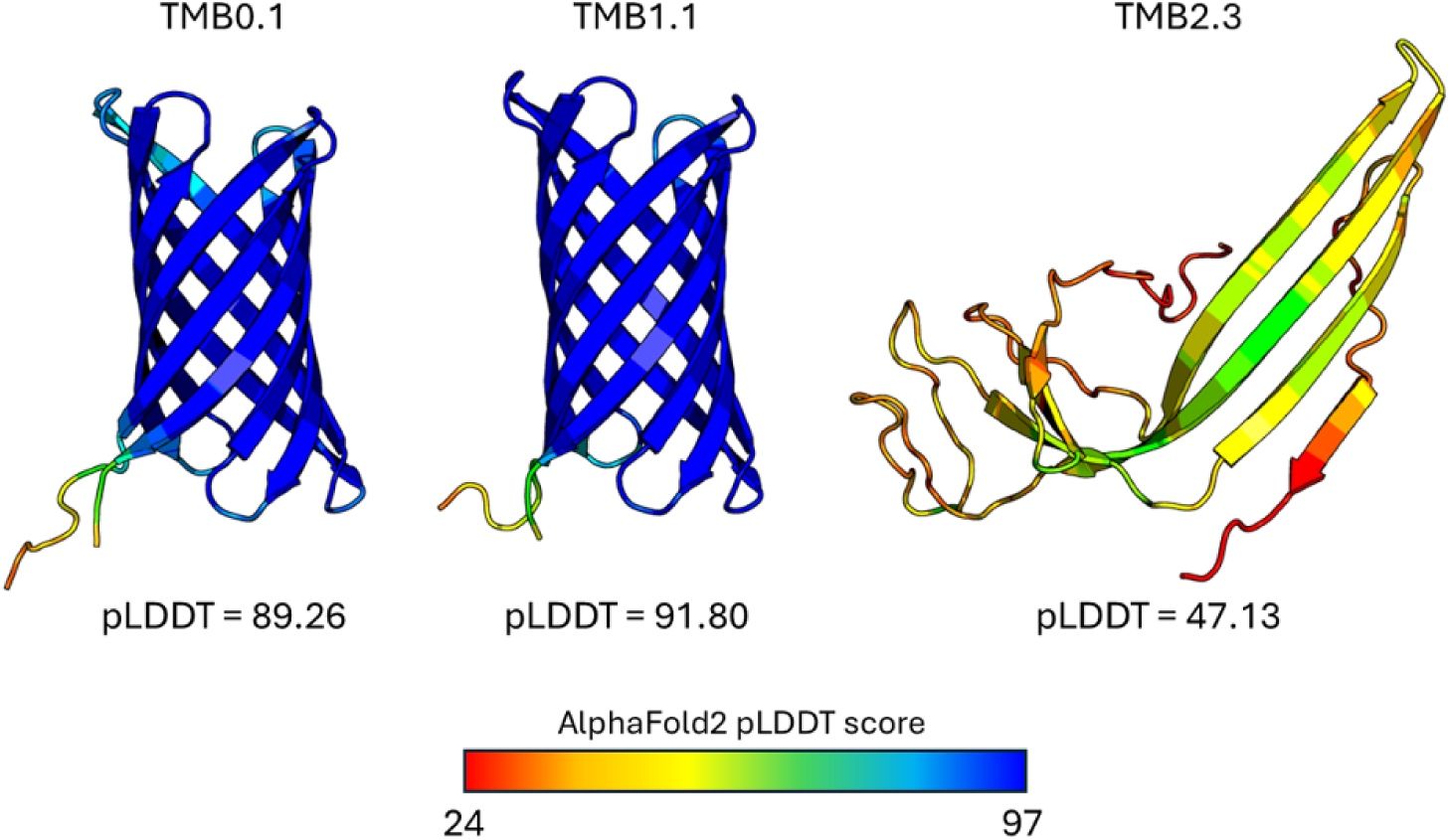
AlphaFold2 predictions of the structures of synthetic sequences TMB0.1, TMB1.1, and TMB2.3. Sequences TMB0.1 and TMB1.1 (which aggregate) are predicted by AlphaFold2 to form stable transmembrane β-barrel structures with high confidence (pLDDT confidence scores >89.0). By contrast, the predicted structure of TMB2.3, which successfully folded in vitro, is an incomplete barrel that is attributed a low confidence score by AlphaFold2. The secondary and tertiary structures of this design are more weakly encoded into the sequence, likely because of the use of negative design. AlphaFold2 was run using ColabFold in single sequence mode; 3 recycles and energy minimization with the AMBER force field.

**Figure S6.**
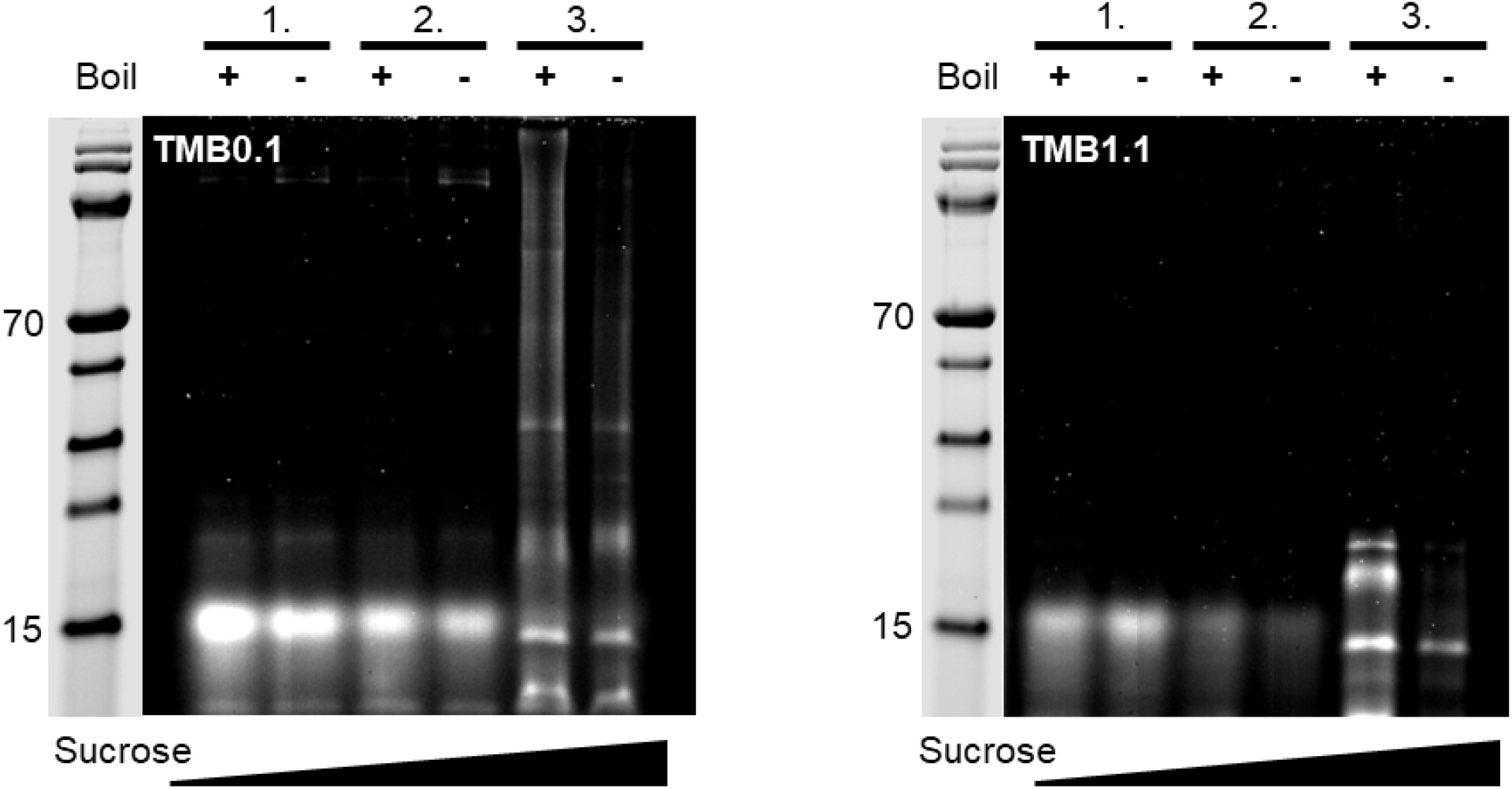
Partitioning profiles of TMB0.1 (left) and TMB1.1 (right) after cell-free expression and sucrose density gradient ultracentrifugation in the presence of E. coli Polar lipids LUVs. Fraction (1.) denotes the top fractions containing the LUVs which floated to the interface between 0% and 30% sucrose solutions. Fraction (2.) contains soluble proteins that are not associated with LUVs. Fraction (3.) contains insoluble and aggregated proteins and components of the cell-free expression kit.

**Figure S7.**
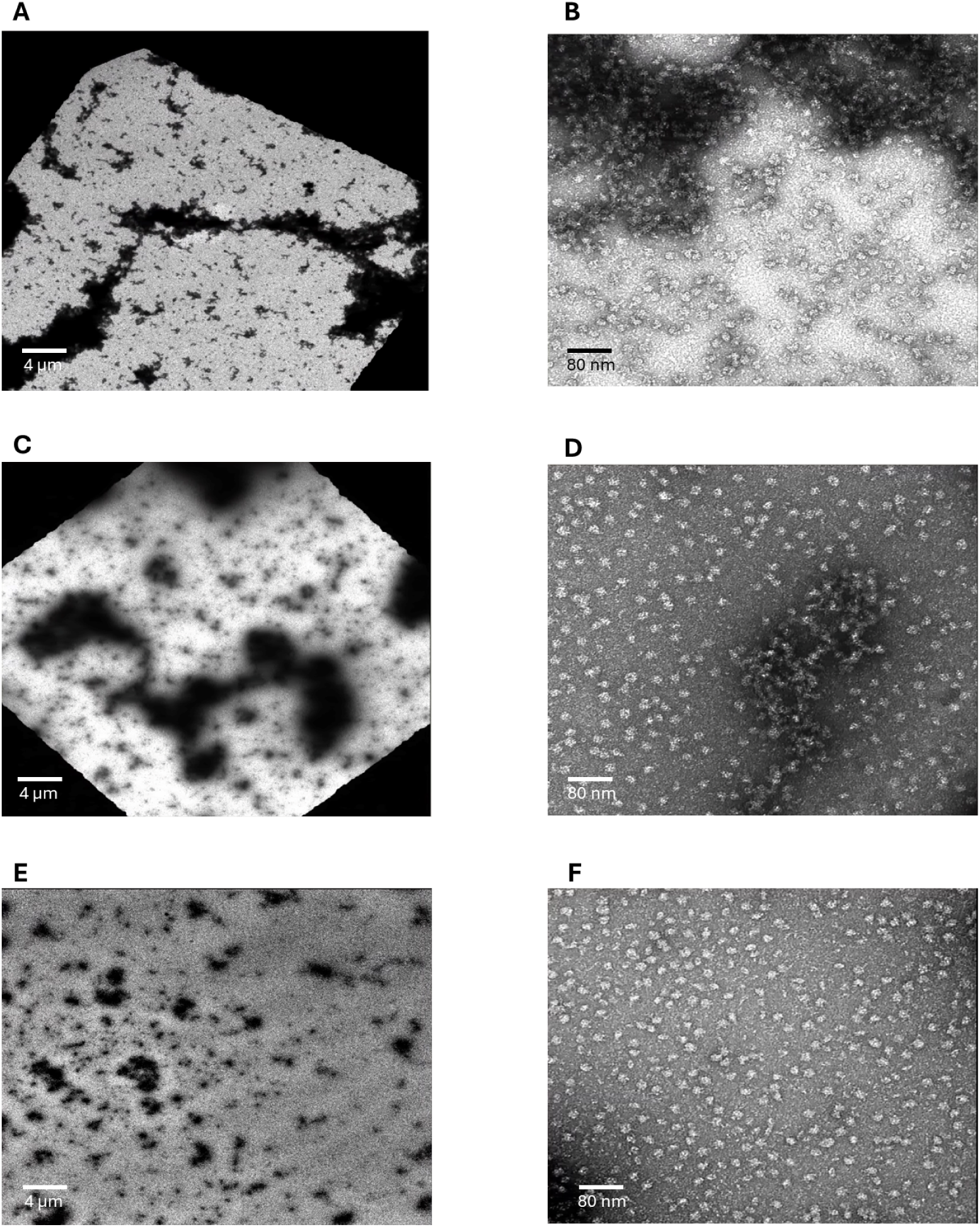
nsEM visualization of TMB0.1 (A, B), TMB1.1 (C, D) and TMB2.3 (E, F) at 2,000 (left) and 60,000 (right) nominal magnification. In the case of TMB0.1 and TMB1.1 aggregates formed by clustered ribosomes are clearly visible at low and high magnification. In case of TMB2.3 the ribosomes are visible as single particles at high magnification, whereas at low magnification only small patches of clustered ribosomes are observed, indicating low aggregation propensity.

**Figure S8:**
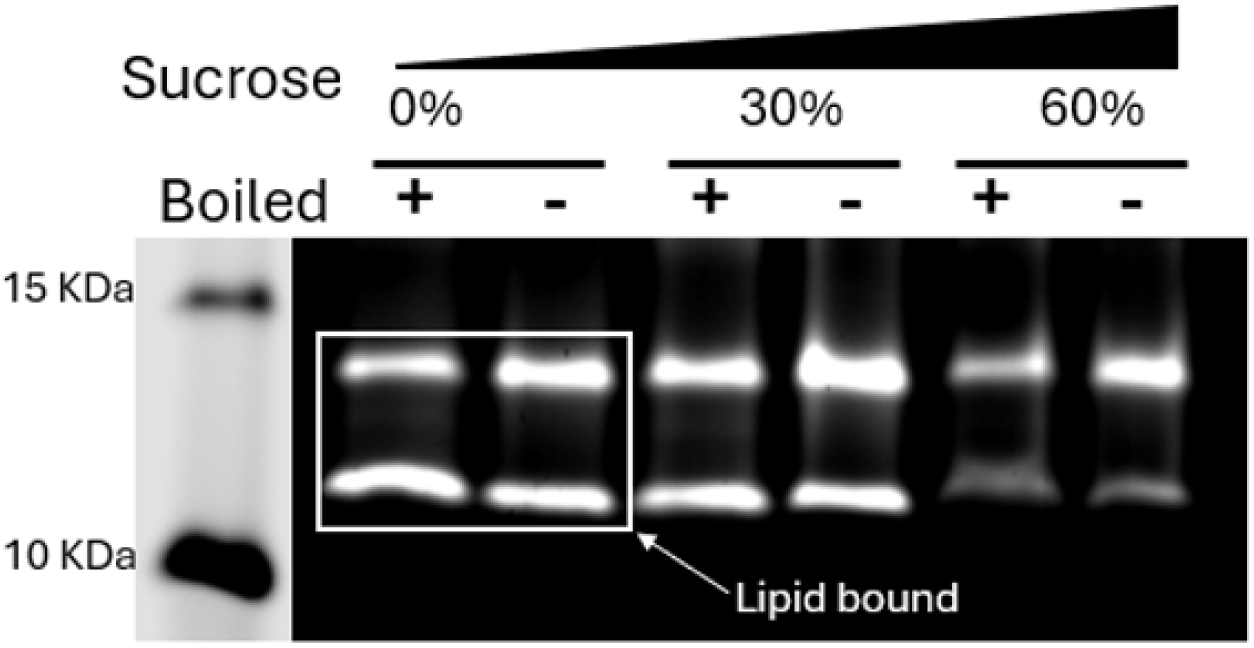
TMB2.17 is distributed between the lipid-bound, water-soluble and aggregated fractions after IVTT and sucrose gradient ultracentrifugation.

**Figure S9.**
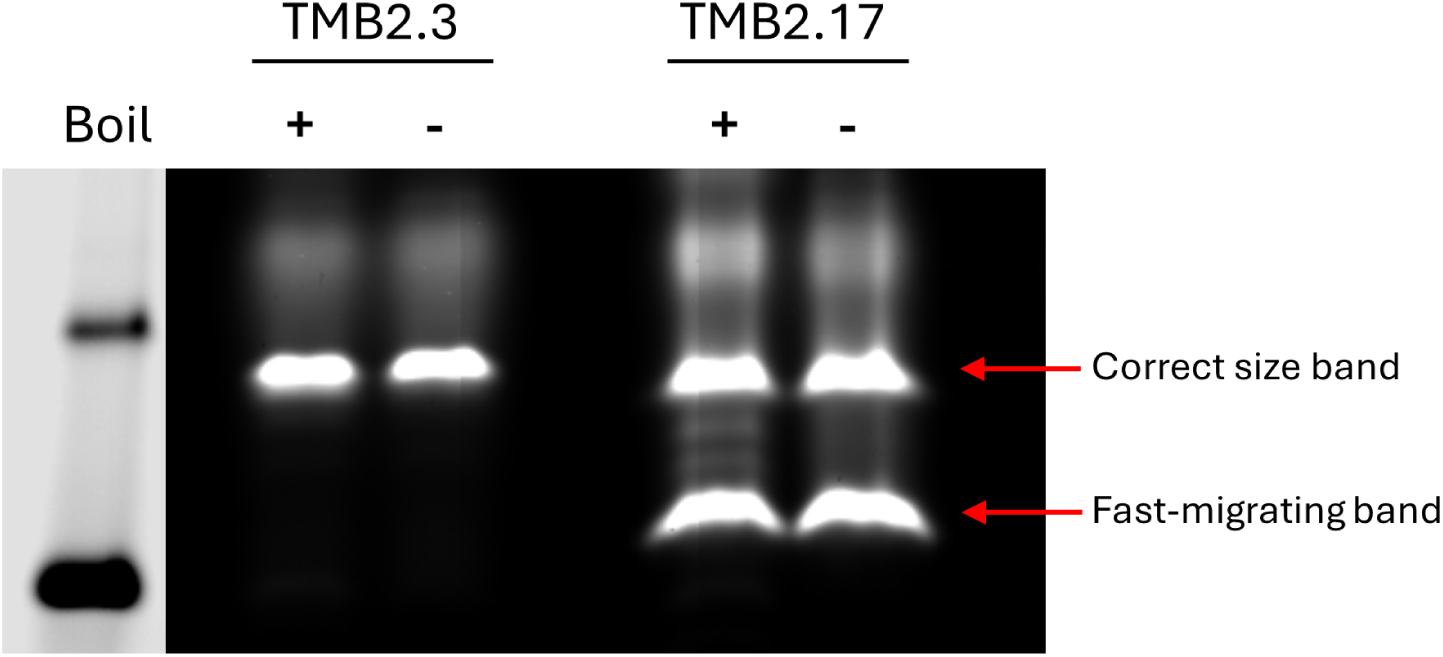
SDS-PAGE analysis of designs TMB2.3 and TMB2.17 expressed in the cell-free system in the presence of DUPC LUVs. TMB2.17 consistently showed a fast-migrating band resistant to boiling, which is not present in TMB2.3.

**Figure S10:**
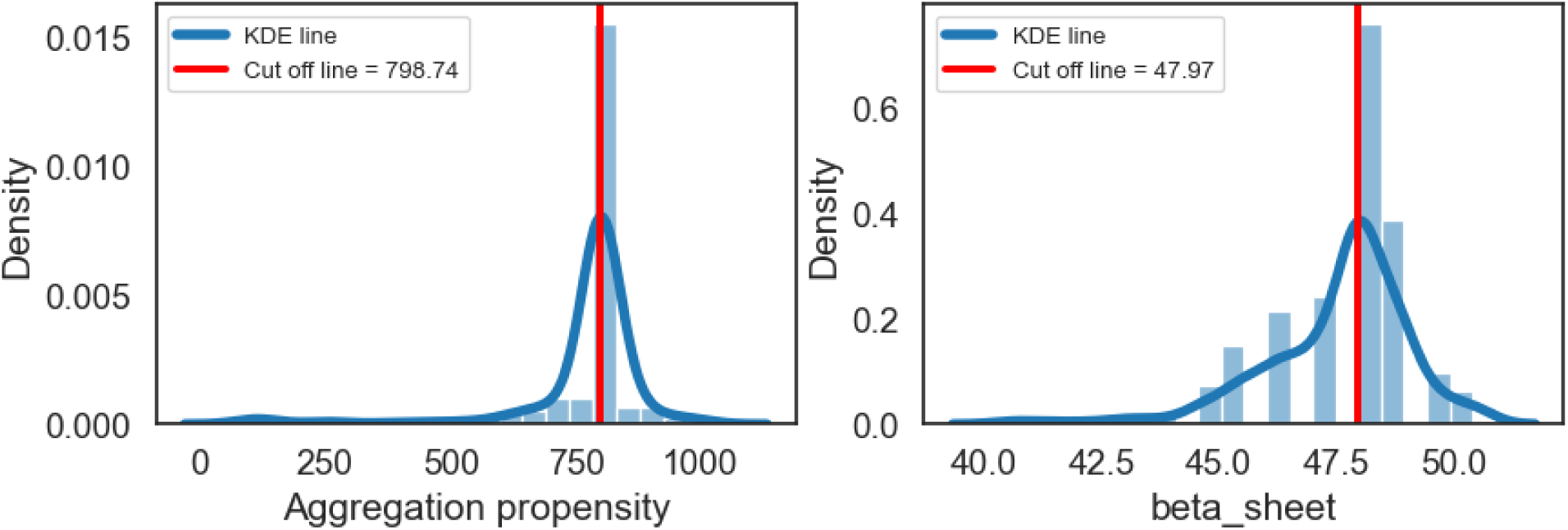
Distributions of β-aggregation scores (predicted with Tango, left) and the fraction of the polypeptide that is assigned β-sheet secondary structure by RaptorX (right), for all substitutions in the first β-strand of TMB2.17 that were predicted to have a stabilizing, neutral, or moderately destabilizing effect on protein stability. The ΔΔG upper boundary energy score was fixed at 3.0 Rosetta Energy Units. The red bar indicates the values of these metrics computed for the wild-type TMB2.17 sequence. While some variants score better than the wild-type on some metrics, no variant was found to score better on all the imposed metrics.

**Figure S11:**
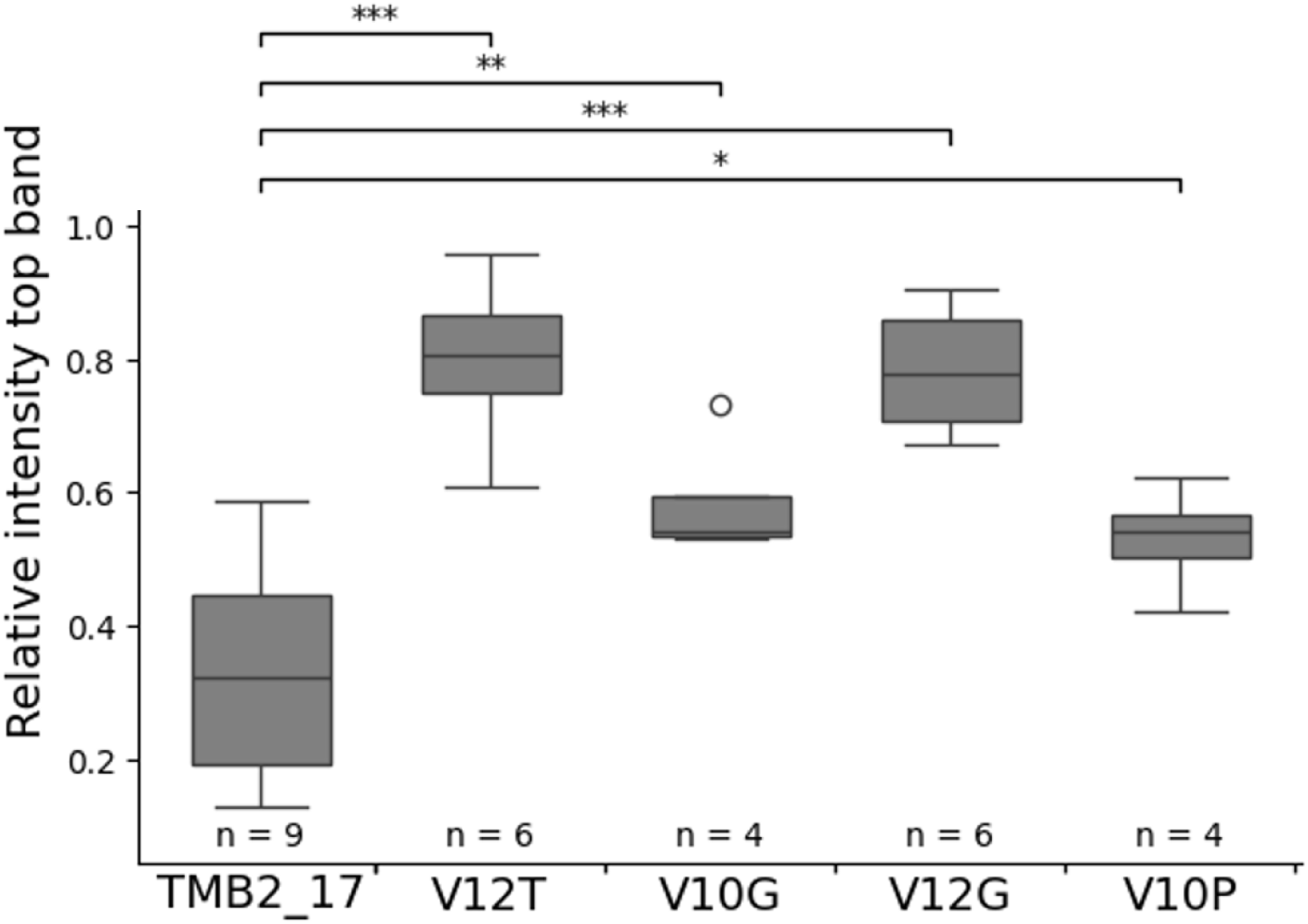
Relative intensity of the major (upper) SDS-PAGE band of TMB2.17; calculated as the intensity of the upper band divided by the sum of intensities of the upper and lower bands. The intensity obtained for the wild-type TMB2.17 is compared to the four folded variants using a Student’s t-test. (*p<0.05; **p<0.01; ***p<0.001).

**Figure S12.**
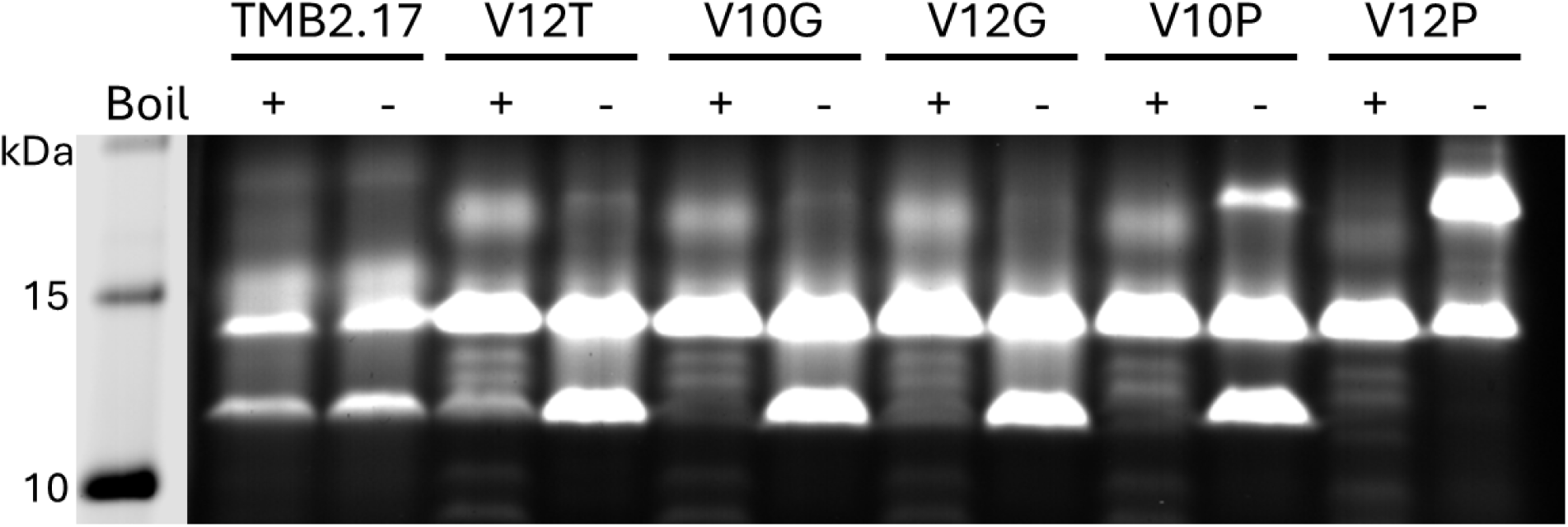
SDS-PAGE of TMB2.17 and its variants. The secondary, fast-migrating band becomes heat-sensitive in the β-strand disruptive variants.

**Figure S13.**
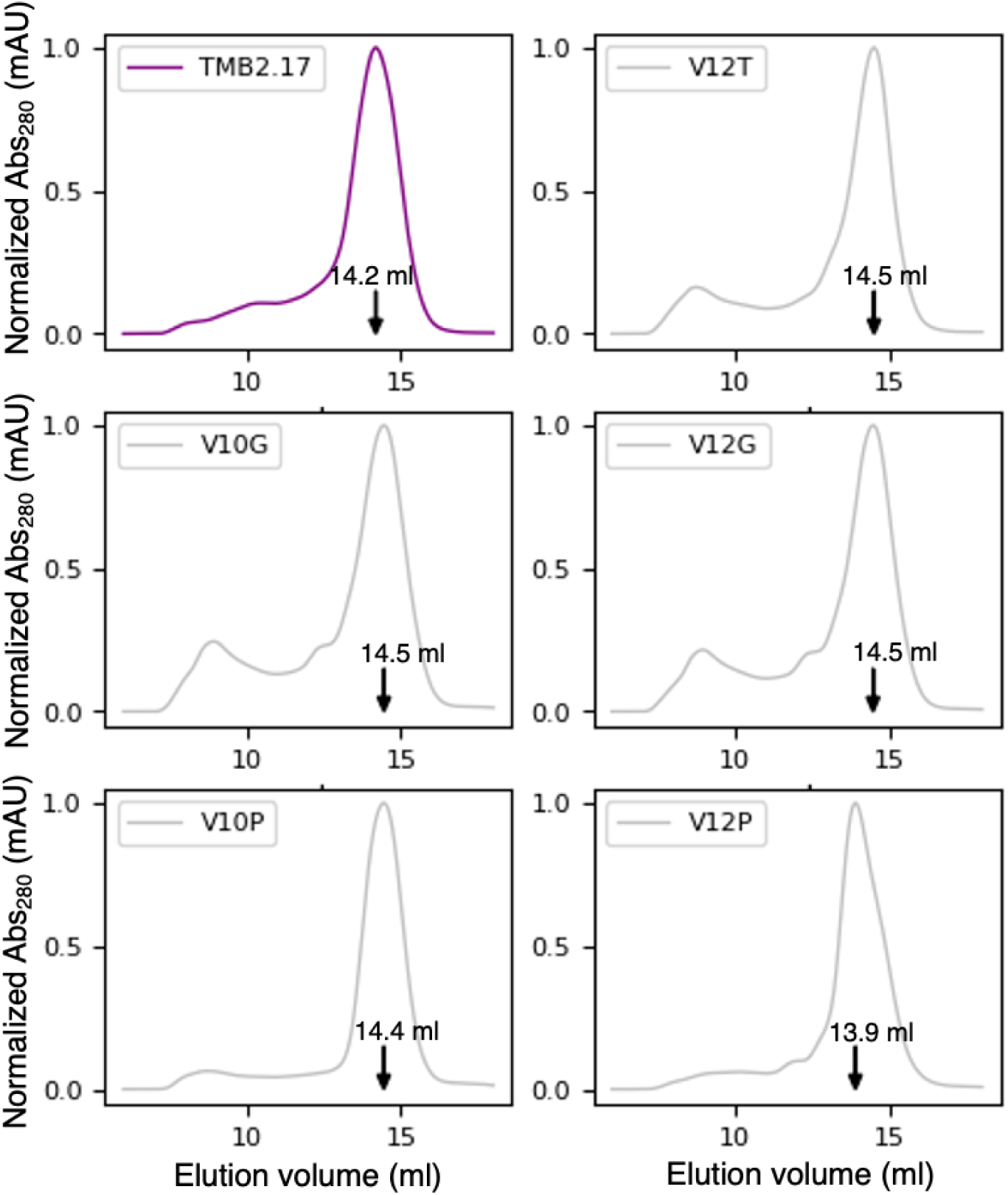
TMB2.17 and its variants elute as major peaks with similar elution volumes in SEC.

**Figure S14:**
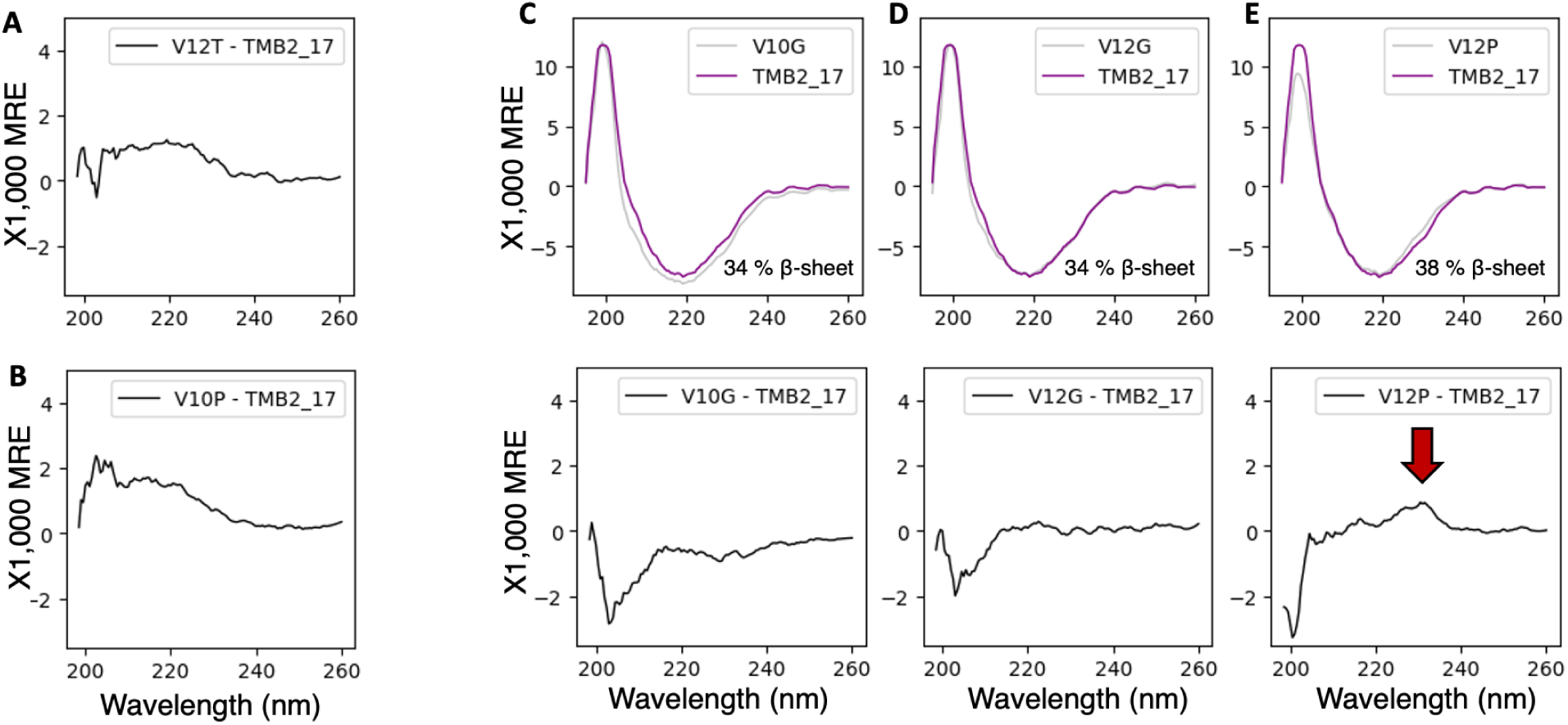
Far UV circular dichroism spectra of TMB2.17 and its variants. (A) Spectral difference between variant V12T and TMB2.17-showing a small increase in stable β-sheet content. (B) Spectral difference between variant V10P and TMB2.17 - showing a small increase in stable β-sheet content. (C) Overlaid spectra of variant V10G (gray) and TMB2.17 (purple) (top) and spectral difference (bottom) - corresponding to a decrease in stable β-sheet. (D) Overlaid spectra of variant V12G (gray) and TMB2.17 (purple) (top) and spectral difference (bottom) - corresponding to a decrease in stable β-sheet. (E) (C) Overlaid spectra of variant V12P (gray) and TMB2.17 (purple) (top) and spectral difference (bottom) - with a marked difference around 230 nm.

**Figure S15.**
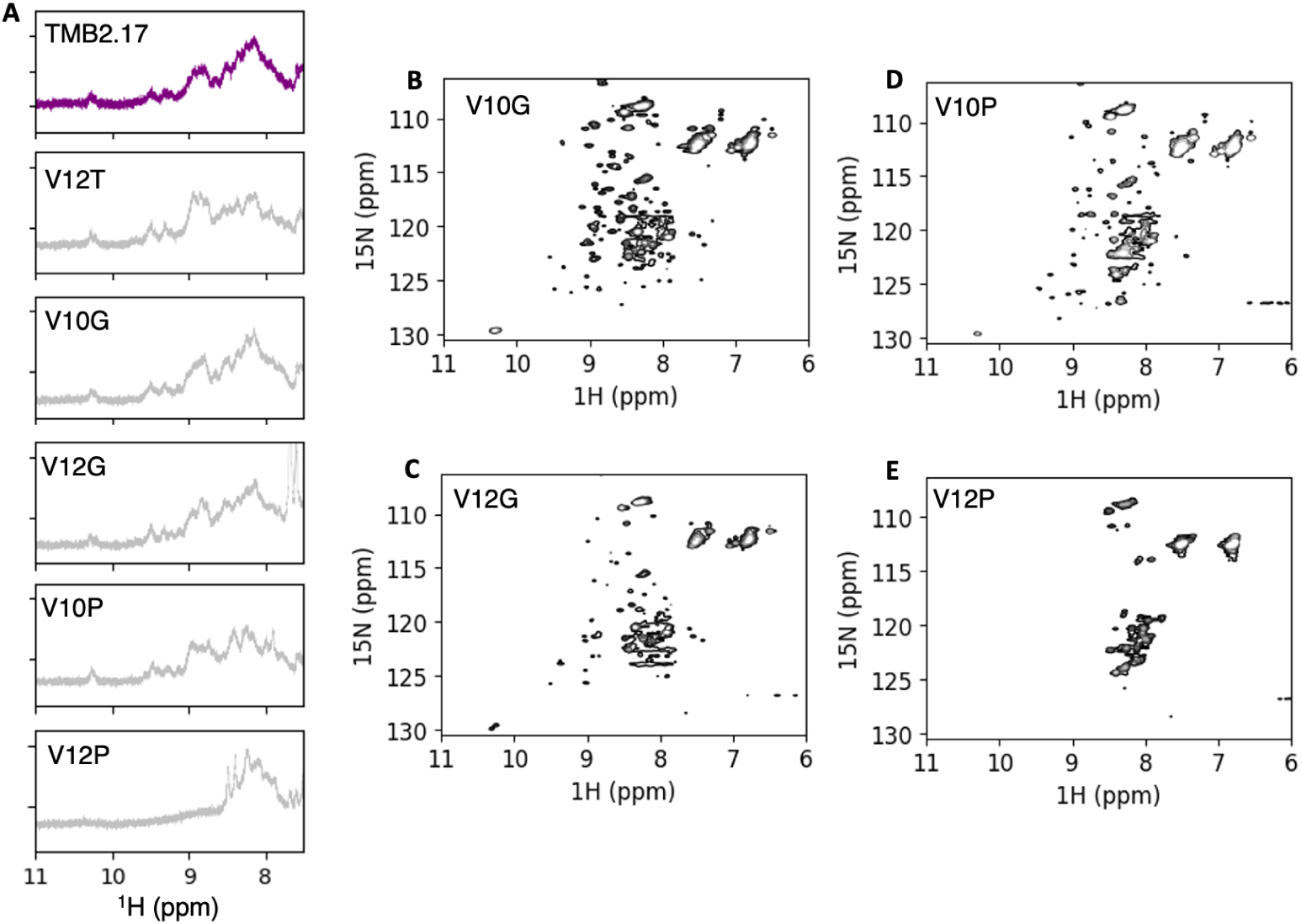
NMR spectra of TMB2.17 and its variants. (A) One-dimensional ^1^H NMR spectra. TMBs spectra are characterized by the presence of distinct peaks between 10.5 and 8.5 ppm. (B-E) ^1^H-^15^N HSQC NMR spectra of variant V10G (B), V12G (C), V10P (D) and V12P (E).

**Figure S16.**
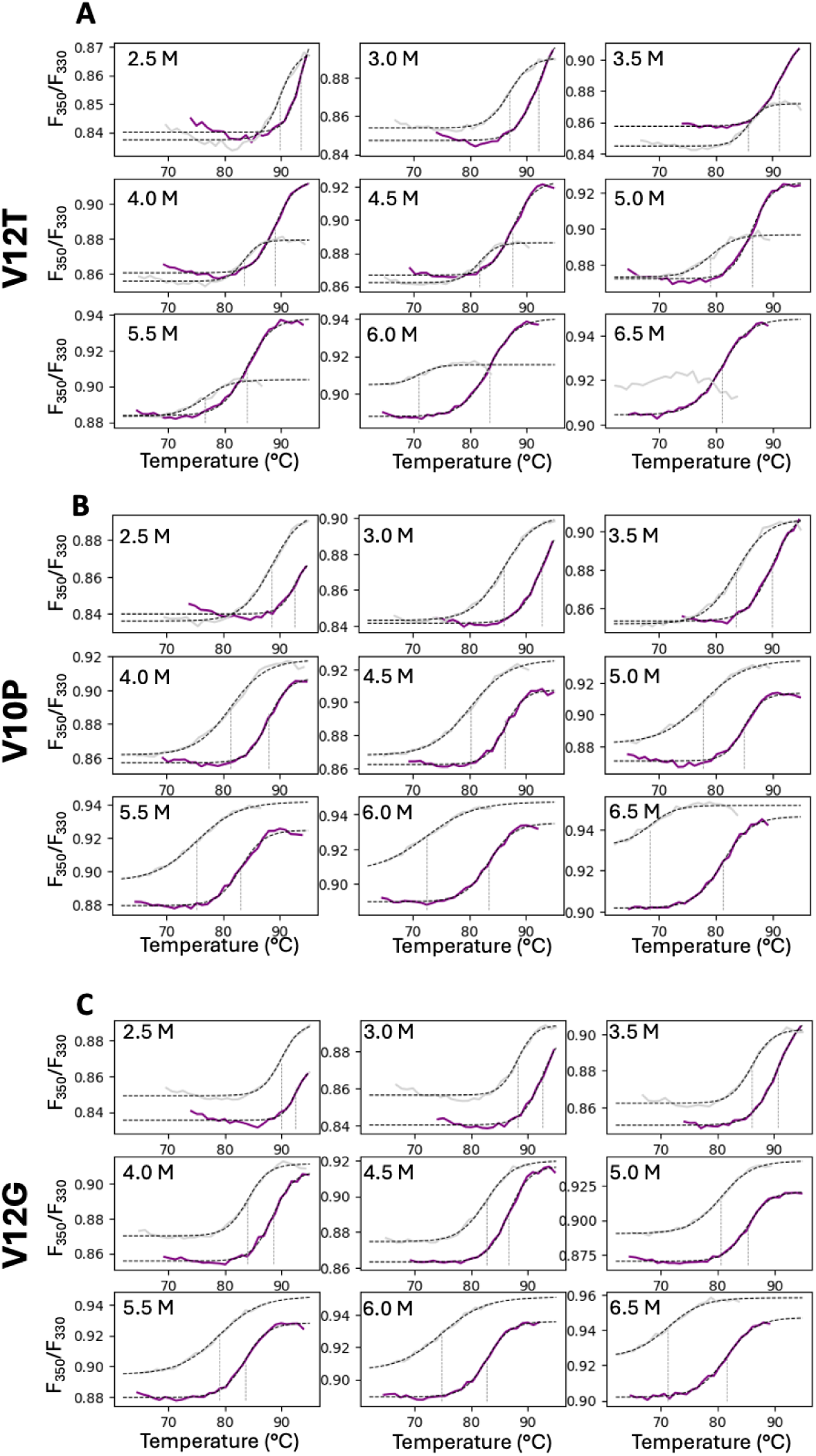
Unfolding transitions of variants V12G, V10P, and V12G in DPC detergent micelles upon temperature increase and in the presence of different concentrations of urea (2.5M-6.5M). The transitions were monitored based on the shift in intrinsic tryptophan fluorescence emission from 330 nm to 350 nm (F_350_/F_350_). The unfolding transitions of the wild-type TMB2.17 are overlaid in purple to each variant (gray) for reference. The melting temperatures (Tm) are indicated with dashed lines. Fluorescence data collected in 7M urea are not shown because TMB2.17 and all the variants were unfolded in those conditions.

## Supplementary tables

**Table S1:**
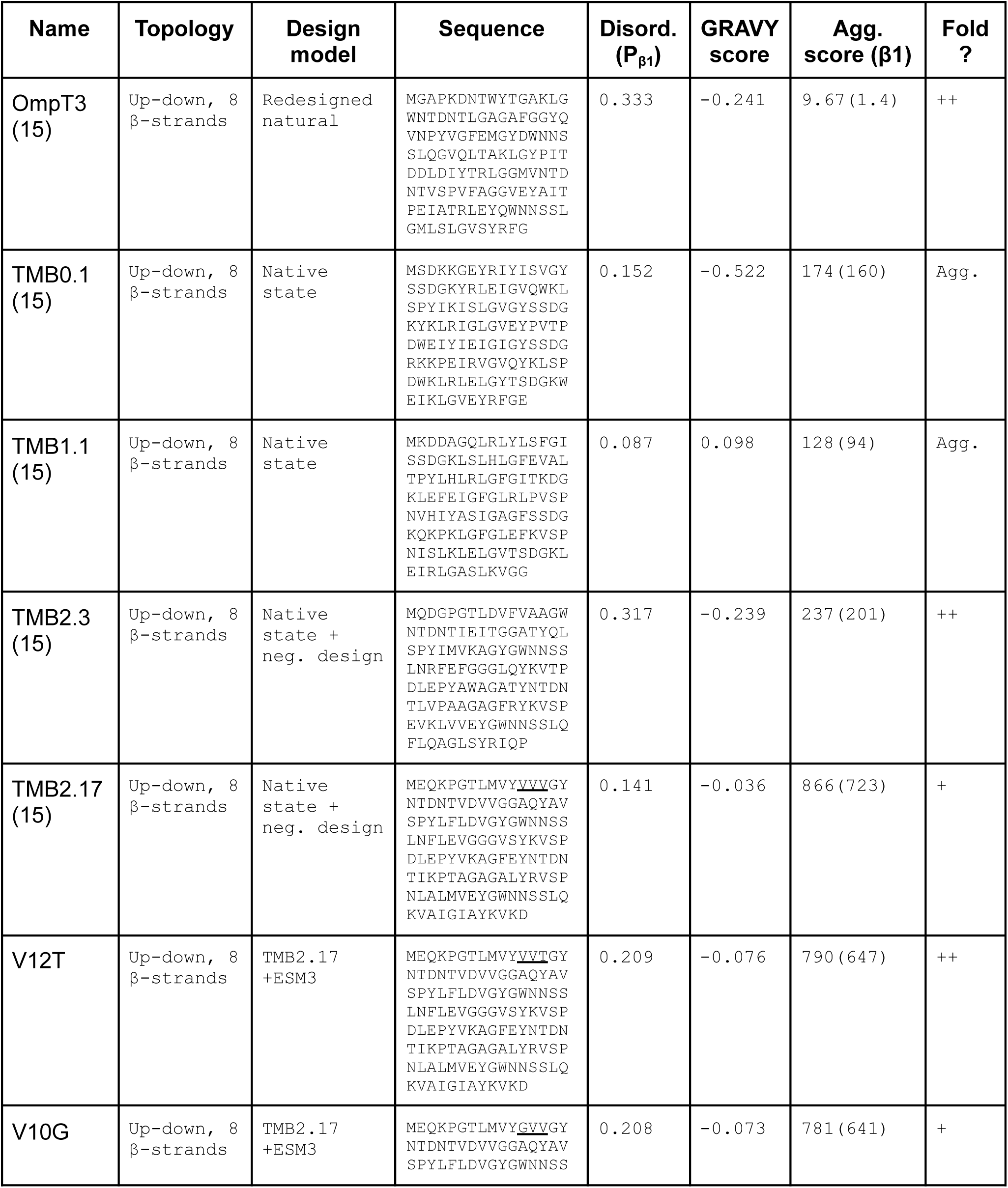
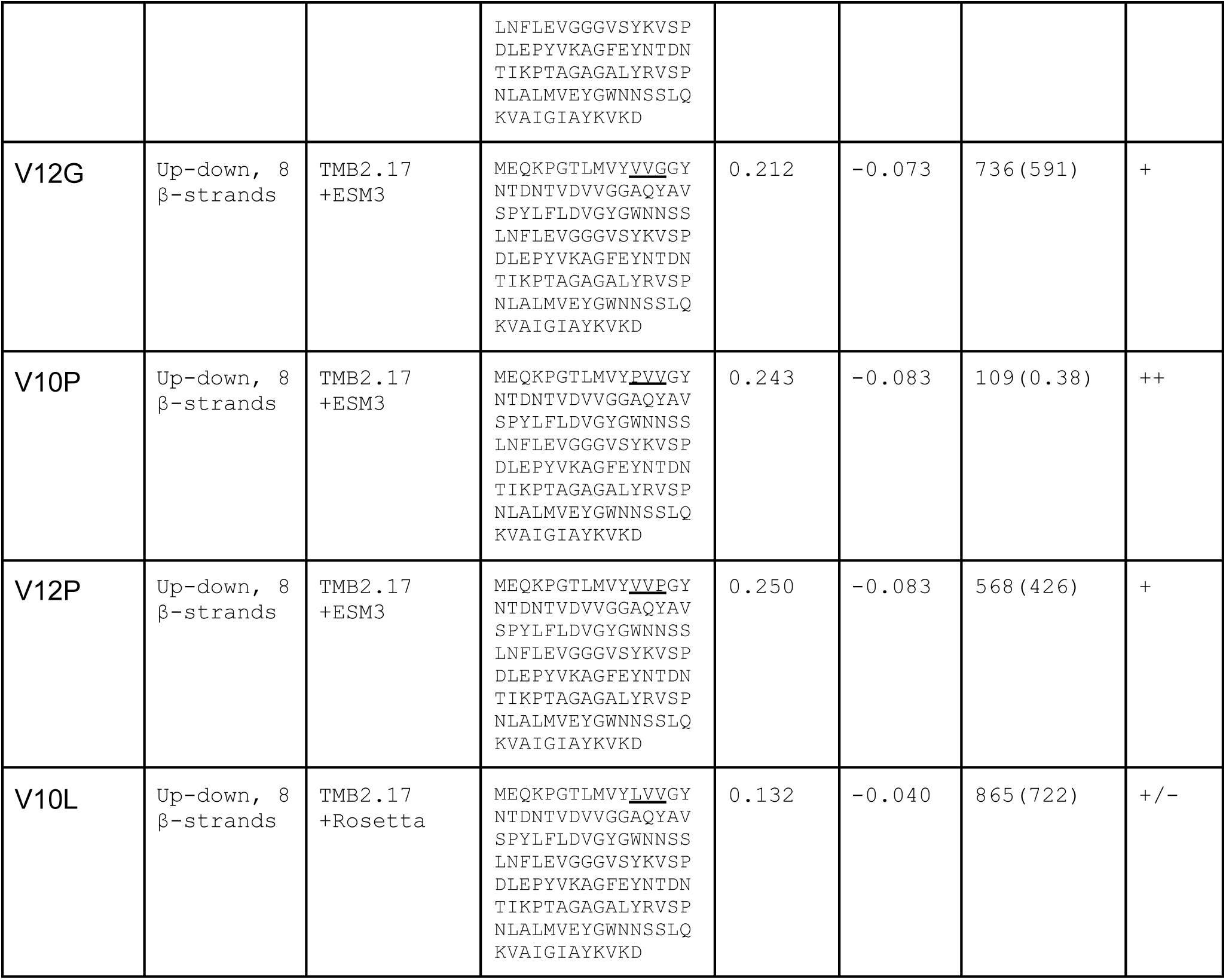
Sequences and properties of the designs used in this study. Local disorder probability was predicted with IUPred3 (short disorder, no smoothing) and averaged across β-strand1 residues. The Aggregation score was predicted using Tango. The total aggregation score for the full-length sequence and the sum over residues belonging to β-strand1 are reported in the table.

**Table S2.**
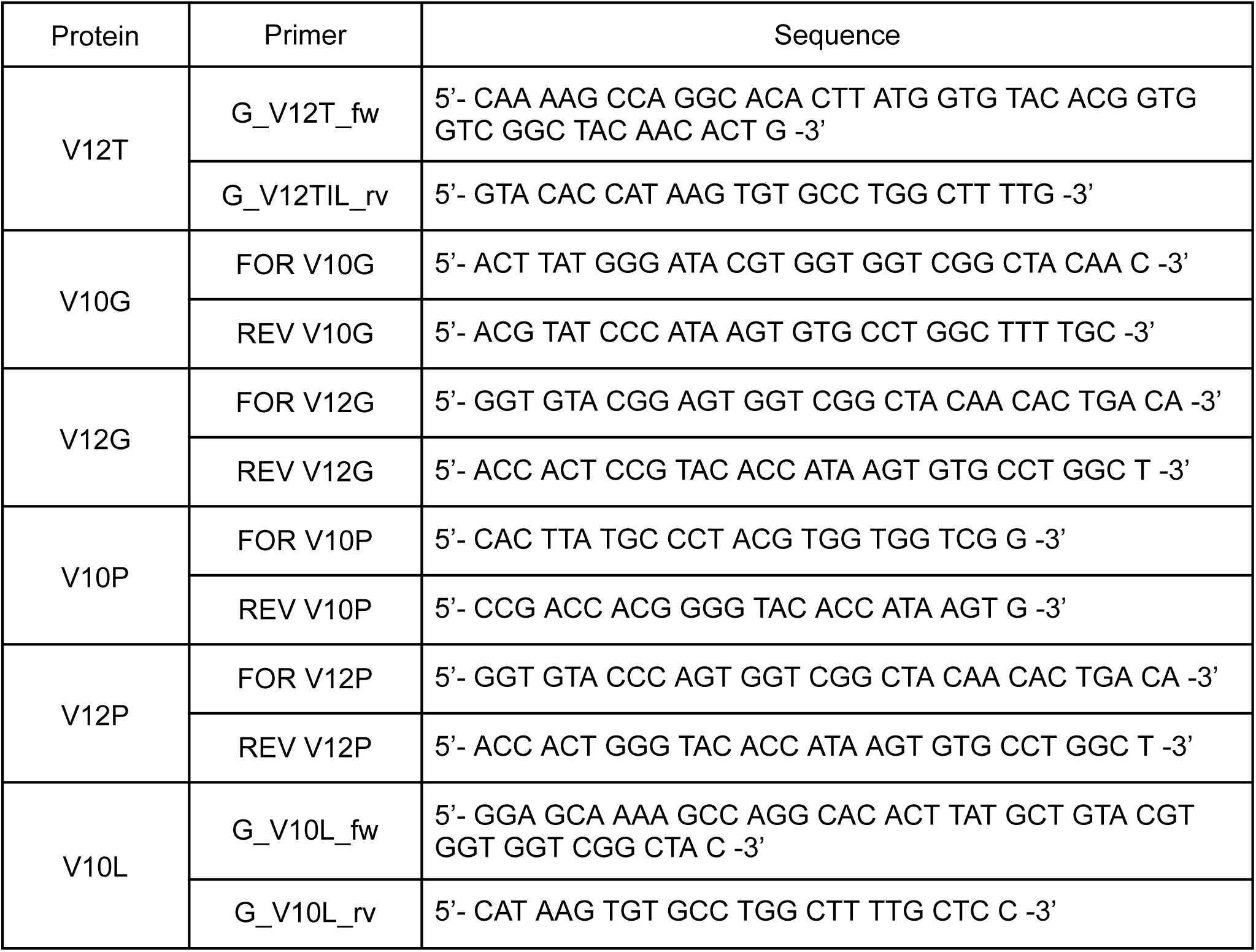
Primer sequences used to perform mutational PCR on TMB2.17.

